# Biological Signaling Pathways Associated with Late Tendon Development in Chicken Embryos and Their Dependence on Mechanical Stimulation

**DOI:** 10.64898/2026.11.02.763759

**Authors:** Maria L. Canonicco-Castro, Maxwell A. Konnaris, Benjamin E. Peterson, Niamh E. Buckley, Rebecca A. Rolfe, Paula Murphy, Helen McCarthy, Nicholas Dunne, Spencer E. Szczesny

**Affiliations:** Department of Biomedical Engineering, The Pennsylvania State University, University Park PA, USA; Program in Bioinformatics and Genomics, The Pennsylvania State University, University Park, Pennsylvania, USA; School of Pharmacy, Queen’s University Belfast, Belfast, Northern Ireland; Zoology, School of Natural Sciences, Trinity College Dublin, Dublin, Ireland; School of Mechanical & Manufacturing Engineering, Dublin City University, Dublin, Ireland; Biodesign Europe, Dublin City University, Dublin 9, Ireland; Advanced Manufacturing Research Centre (I-Form), School of Mechanical and Manufacturing Engineering, Dublin City University, Dublin 9, Ireland; Research Centre for Medical Devices (CÚRAM), Biomedical Sciences, University of Galway, Galway, Ireland; Department of Orthopaedics and Therapy Services, The Pennsylvania State University, Hershey PA, US

**Keywords:** Tendon, Embryonic development, RNA-Sequencing, Mechanotransduction, Paralysis

## Abstract

During embryonic and neonatal development, tendons exhibit rapid changes in mechanics and structure, including increases in collagen content, crosslinking, and fibril length. These functional and structural changes are partly dependent on mechanical stimulation. However, the specific mechanobiological mechanisms driving these changes remain poorly understood. The objective of this study was to identify the potential biological mechanisms and pathways driving tendon maturation into load-bearing materials using an embryonic chick model. Using bulk RNA sequencing, gene set enrichment analysis, and transcription factor analysis, we produced a transcriptomic profile of chick hindlimb tendons during late embryonic development (E16, E18, and E20) and tendons from rigid and flaccid paralysis-treated embryos. Additionally, we assessed the results from two differential expression analysis models to improve the replicability of our findings. Our results suggest that the largest transcriptomic response occurs between embryonic day 16 and 20, and that among paralysis conditions, rigid paralysis produces the greatest effect relative to controls. Across development, we found positive enrichment in processes related to mechanotransduction, extracellular matrix remodeling, cell adhesion, and contractility, and identified candidate pathways upregulated with development, including TGF-β, PI3K-AKT-mTOR, Hippo, circadian rhythm, and focal adhesion signaling. In paralysis-treated embryos, our data suggest impairment in extracellular matrix remodeling and mechanotransduction events. Specifically, we found that both modes of paralysis are associated with a cellular stress response and downregulation of WNT signaling. Together, our findings identify candidate mechanotransduction pathways that are associated with late tendon maturation and altered by paralysis, which may inform tendon tissue engineering and regenerative approaches.

## 1. Introduction

Tendons and ligaments are connective tissues that enable proper joint movement and locomotion. Unfortunately, they are commonly injured and account for approximately 36% of all nonfatal occupational injuries in the United States annually^1^. Following injury, tendons and ligaments heal through a scar-mediated process that produces a disorganized extracellular matrix and mechanically inferior tissue, increasing the risk of re-injury and re-rupture^2–5^. Certain tendon and ligament ruptures (e.g., anterior cruciate ligament, chronic Achilles tendon tears, supraspinatus tendon) may require reconstruction via autologous or allogeneic grafts^6–8^; however, these approaches are limited by donor site morbidity and inferior outcomes, respectively^9^. These limitations have driven interest in the development of tissue-engineered tendon/ligament replacements. Although advances have been made, no engineered construct has successfully replicated the structural organization and long-term load-bearing capacity of native tendons or ligaments^10,11^. A fundamental barrier to the successful creation of a tendon/ligament replacement tissue is that the essential biological mechanisms that establish tendon/ligament structure and function during development are still unclear^12^. Therefore, further study of these structural, functional, and biological changes that drive tendon/ligament development is critical for advancing tissue engineering and regenerative medicine strategies.

The structural and mechanical changes that occur during tendon development have been well characterized. During early development, cells align into columns through cadherin-11 mediated cell-cell junctions^13^ and begin depositing short collagen fibril segments into longitudinal intracellular channels to form the initial collagenous extracellular matrix^14–16^. At later stages of tendon development, there is a decrease in the cell volume fraction, reflecting a transition from a cell dense tissue to one composed predominantly of extracellular matrix^17–19^. Specifically, at embryonic day 16 (E16) in the chick and post-natal day 10 (P10) in mice, there are large increases in collagen content, fibril elongation, and intrafibrillar crosslinking^14,15,20^ as well as a rapid increase in tissue mechanical properties^16,21,22^. Recent work from our lab has demonstrated that fibril elongation and increased collagen crosslinking are the main drivers of the elevated mechanical properties that are observed during late tendon development^23^. Importantly, this maturation process is dependent on mechanical stimulation from muscle contractions with paralysis leading to reduced mechanical stiffness, strength, and collagen fiber alignment^19,23–25^. Together, these findings define the structural and mechanical changes that occur during tendon development and suggest that mechanobiology plays a central role in tendon maturation. However, the specific biological signaling pathways mediating the structural changes that occur during late tendon development are unclear.

Possible candidates are the transcription factors and pathways that are important for early tendon development. At the molecular level, the transcription factor Scleraxis (Scx) is one of the earliest markers of tendon progenitor cells and plays a central role in tenogenic differentiation^26,27^. Additional transcription factors, including Mohawk (Mkx) and early growth response proteins (Egr1, Egr2) contribute to cellular differentiation, collagen production, and matrix maturation^28–31^. Furthermore, signaling pathways such as transforming growth factor-β (Tgf-β) and fibroblast growth factor (Fgf) have been identified as key promoters of tendon fate specification and mediators of mechanotransduction during early development^32,33^. In contrast, Wnt/β-catenin signaling has been shown to repress tenogenic gene expression *in vivo* and suppress *Scx* and *Mkx* expression *in vitro*^34,35^.

Contrary to early development, the molecular mechanisms driving late-stage tendon maturation remain less understood. Given the mechanosensitive nature of this process, recent studies have begun to identify candidate pathways involved in late development. For example, Yap/Taz signaling has been implicated in late tendon maturation, as it is responsive to embryonic movement^19^. Transcriptomic evidence further supports a role for Yap signaling with Yap target gene expression observed in embryonic chick tendons between E11 and E17^36^. However, while conditional loss of *Yap1* at post-natal stages in mice impacted collagen expression and matrix organization, there was no effect on the ultrastructural or mechanical properties of tendons^37^. Similarly, focal adhesion kinase (Fak) signaling affects tendon size and cellular composition during post-natal growth in mice^38^; however, *Fak* knockout did not produce a decrease in the tendon material properties^38^. Therefore, while prior work suggests that mechanosensitive signaling pathways contribute to tendon maturation^23–25^, the specific pathways driving the substantial changes in structure and mechanics observed during late tendon development are still unclear. Additionally, the transcriptomic differences between normally developing and mechanically perturbed (i.e., paralyzed) embryos have yet to be defined, and it is still unclear whether these mechanosensitive pathways are conserved across species. Therefore, despite these advances, there are still significant gaps in our understanding of late-stage tendon development.

The objective of this study was to identify the potential mechanobiological mechanisms associated with tendon maturation and mechanical perturbation using a chick embryo model of late tendon development. To accomplish this, we performed bulk RNA sequencing of the hindlimb tendons in normally developing embryos (E16, E18, and E20) as well as in embryos subjected to rigid and flaccid paralysis. We then applied two analytical methods (DESeq2 and ALDEx3) to measure changes in gene expression. Given its common use in the literature, DESeq2 facilitates the comparison of our results with prior findings, while ALDEx3 provides potentially more reproducible results by incorporating uncertainty regarding differences in total gene expression between sample groups (i.e timepoints or conditions)^39,40^. This also enables the identification of pathways that are dependent on a particularly analytical approach versus those that are more robust findings in our dataset. Finally, we performed gene set enrichment analysis and transcription factor overrepresentation analysis to identify the biological processes, signaling pathways, and transcriptional regulators underlying normal tendon maturation and their dependence on mechanical stimulation.

## 2. Materials and Methods

### 2.1 Sample Collection and Treatments

Fertilized White Leghorn eggs were obtained from the Penn State Poultry Education and Research Center and incubated at 37.7°C from embryonic day 0 (E0). At E3, eggs were windowed following standard protocols^25,42^. For normal development samples, windowed embryos were sacrificed by decapitation at E16, E18, and E20, which span the rapid changes in tendon structure and mechanics observed in late tendon development^21,23^. For paralysis samples, embryos were treated by dropwise administration of decamethonium bromide (DMB) or pancuronium bromide (PB) onto the chorioallantoic membrane. DMB induces a “rigid” paralysis through the irreversible binding of acetylcholine receptors at the post-junctional membrane of the motor end plates producing a contracture-like paralysis^43–47^. PB induces a “flaccid” paralysis by competitively binding to the acetylcholine receptors, impairing downstream muscle activation and off-loading both the muscle and tendon tissue^45,47^. Treatment began at E13 with an initial dose of 100 µL of 0.2% DMB (TCI America) or 0.2% PB (MP Biomedicals) in sterile HBSS with 1% antibiotic-antimycotic (AA), followed by daily 50 µL doses from E14 through E19. Vehicle controls received sterile HBSS with 1% AA at volumes matched to the treatment groups. This treatment regimen is sufficient to reduce embryonic motility through E20^25^. All paralysis treatment groups and vehicle controls were sacrificed by decapitation at E20.

For all samples, *flexor digitorum longus* (FDL) and *flexor digitorum brevis* (FDB) digit II tendons were harvested from the hindlimbs. Each biological replicate (n=3 for normal development, n=4 for paralysis) consisted of eight pooled tendons (two FDL and two FDB tendons harvested from the hindlimbs of two embryos). All embryos were staged according to Hamburger and Hamilton (HH) criteria^48^, using the third toe length to confirm developmental stage at E16 (HH42), E18 (HH44), and E20 (HH46). Normal development samples were flash-frozen in liquid nitrogen (LN), stored at −80°C, and shipped on dry ice to Genewiz (Azenta Life Sciences) for analysis. Paralysis and vehicle control samples were placed in TRIzol, flash frozen in LN, and stored at −80°C until RNA extraction and shipment.

### 2.2 RNA Extraction

The frozen paralysis and vehicle control samples were homogenized using polypropylene pestles. RNA was extracted using chloroform and purified using a PureLink RNA Mini Kit (Invitrogen) following manufacturer’s instructions. RNA concentration was quantified using a Qubit 4 Fluorometer (Thermo Fisher Scientific), and RNA integrity numbers (RINs) were determined using a TapeStation (Agilent). For normal development samples, RNA extraction was performed by Genewiz (Azenta Life Sciences) following their standard procedures for frozen tissue. Differential gene expression analysis revealed significant differences between the vehicle control and E20 samples, potentially attributable to differences in RNA extraction, antibiotic treatment, or windowing (Supplementary Fig. 1). To account for these discrepancies, normal development and paralysis samples were not directly compared in downstream analyses.

### 2.3 RNA Sequencing

RNA samples were subjected to quality control (RIN > 6.0, RNA concentration ≥ 50 ng/µL) prior to sequencing. Library preparation and sequencing were performed by Genewiz (Azenta Life Sciences) using poly(A) selection for mRNA enrichment. Libraries were sequenced on an Illumina HiSeq platform (2×150bp paired-end, ∼50 million reads per sample). Raw FASTQ files were assessed for sequencing quality using FastQC, and sample-level quality-control reports were aggregated using MultiQC. Adapter trimming and read-quality filtering were performed using fastp. Trimmed reads were aligned to the *Gallus gallus* reference genome using STAR, and transcript abundance was quantified using Salmon. The genome index and transcript annotation were generated from the *Gallus gallus* assembly bGalGal1.mat.broiler.GRCg7b with the corresponding Ensembl release 115 GTF annotation file. Salmon-derived gene-level count tables were used as input for downstream analyses, and sample-level metadata were constructed from sample names to define the experimental condition for each sample.

### 2.4 Differential Gene Expression Analysis

Sequencing measures only relative expression and provides no information regarding the changes in total gene expression between conditions (i.e., scale)^39^. Consequently, inference about absolute expression, including differential absolute expression, relies on how the unobserved scale is considered. Widely used normalization-based differential-expression tools, including DESeq2, implicitly fix the scale without accounting for uncertainty^40,49^. This has substantial consequences: in benchmark analyses, normalization-based methods that implicitly assume the scale and ignore uncertainty had average false discovery rates exceeding 75%^40,41^. Rather than a fixed assumption of scale, relaxing total-sum scaling (TSS) -- a normalization which assumes the total gene expression is unchanged between conditions -- by explicitly modeling the uncertainty in scale can dramatically reduce false discovery rates and increase sensitivity^40,41^. Therefore, we analyzed the sequencing data with ALDEx3 to potentially improve the replicability of our findings by accounting for scale uncertainty^50,51^ as well as with DESeq2 to facilitate comparison with other bulk RNA-seq analyses given its common use in the literature^52^.

Separate differential analyses were performed for the paralysis and developmental datasets. For the paralysis dataset, the condition factor included CTRL, DMB, and PB, with CTRL as the reference level. For the developmental dataset, it included E16, E18, and E20, with E16 as the reference. Prior to model fitting, low-expression genes were removed by a group-aware filtering procedure: counts per million (CPM) were calculated from the raw count matrix, and genes were retained if they had CPM ≥ 0.5 in at least two samples within at least one experimental group and a total count ≥ 10 across all samples. Genes with zero total counts after filtering were also removed.

To account for scale uncertainty, we fit ALDEx3 using a TSS scale model with γ =0.3 on the log2 scale. Following published guidance^53^, this setting allows modest between-condition differences in total gene expression rather than fixing that difference at zero, effectively adding a conservative margin of approximately 0.5 log2 -fold-change (about 1.4-fold) before significance is declared. Log2 fold-change effect estimates and standard errors were summarized as the median across 2000 Monte Carlo samples. Second, we performed a conventional DESeq2 analysis with adaptive log2 fold-change shrinkage^52^, providing results directly comparable to prior studies in this field. For both methods, results were extracted for each paired contrast (E20 vs E16, E18 vs E16, DMB vs CTRL, PB vs CTRL), and differentially expressed genes (DEGs) were considered significant at a Benjamini-Hochberg adjusted p-value < 0.05 and an absolute log2 fold-change > 1. The effect of relaxing the significance threshold to an adjusted p-value < 0.1 on the number of DEGs identified by each model was also evaluated (Supplementary Fig. 2).

Sample-level structure was evaluated using PCA and clustered sample-correlation heatmaps using ALDEx3-estimated relative abundances (Supplementary Figs. 3 and 4). Specifically, the ALDEx3 log-composition array was summarized by taking the median across Monte Carlo samples for each gene and sample, producing a gene-by-sample matrix of estimated relative log-abundances. These summaries were used for exploratory visualization of sample structure only, with all inference performed on the full Monte Carlo samples. PCA was performed on the transposed matrix after selecting the top 2000 most variable genes. Sample-correlation heatmaps were generated by computing Pearson correlations between samples using the same ALDEx3-derived relative-abundance matrix, followed by hierarchical clustering using one minus the sample-correlation matrix as the distance measure.

### 2.5 Gene Set Enrichment Analysis (GSEA)

For each paired contrast, all genes that passed quality control described above (regardless of p-value or log2 fold-change) were converted to human orthologs using *g:Profiler*^54^ and filtered for genes that had a recognized human gene symbol. When a chicken gene mapped to multiple human ortholog candidates, the original gene symbol was preserved when applicable. Duplicate gene entries were filtered to keep the largest absolute fold change per gene. Through this process, we identified human orthologs for ∼75% of our original genes. For each analysis technique (i.e., DESeq2 or ALDEx3), the genes were ranked by the log2 fold-change, and GSEA^55^ was performed using the *fgsea* package (v1.36.2) in R against four pathway databases (KEGG Legacy, WikiPathways, BioCarta and Reactome (MSigDB c2.cp.v2025.1)) as well as Gene Ontology Biological Processes (GO:BP; MSigDB c5.go.bp.v2025.1). Pathways were included if the gene set contained at least 10 genes identified by our analyses and if at least 5 of these genes were annotated. Statistical significance is displayed in the figures (Fig. 3-6) at adjusted p-value thresholds of 0.05 and 0.1 and a normalized enrichment score (NES) ≥ |1|. To reduce the size of the figures, the pathways and terms were manually curated to those deemed relevant to tendon development. Microsoft Copilot was used to confirm that all the same pathways were considered across all developmental and paralysis datasets; all outputs were verified, and final biological selection was performed by the authors.

### 2.6 Transcription Factor Analysis

Significant orthologous DEGs (log2 fold-change > |1|, adjusted p-value < 0.05) were separated into upregulated and downregulated gene sets and submitted independently to ChEA3^56^ for transcription factor analysis. The integrated TopRank scoring method was used to identify transcription factors most likely to regulate each input gene set, with lower scores indicating greater overlap between a transcription factor’s known targets and the input gene set. Figures include only transcription factors with an integrated score below 0.05.

## 3 Results

### 3.1 Comparison of Differential Gene Expression Between ALDEx3 and DESeq2

As detailed in section 2.4, we used both DESeq2 and ALDEx3 to evaluate differences in gene expression with development and paralysis. This is because, unlike DESeq2, ALDEx3 explicitly models potential differences in mean total gene expression between groups (i.e., scale) rather than making implicit assumptions about this unobserved quantity, which can substantially reduce false-positive rates^40,41^. Furthermore, comparison of the results between each analytical approach helps to identify the findings that remain robust despite this uncertainty. To this end, we compared the number of DEGs identified by each technique and the estimated change in expression. For normal development, across all three comparisons (E18 vs E16, E20 vs E18, and E20 vs E16), DESeq2 consistently identified a greater number of significant DEGs than ALDEx3 (Fig. 1A-C). This difference was most pronounced in the E18 vs. E16 comparison, where ALDEx3 identified only 6 total DEGs (2 unique, 4 shared, 2.4% overlap) compared to 166 by DESeq2 (121 upregulated, 45 downregulated) (Fig. 1A, Supplementary Fig. 5A). The E20 vs. E18 comparison was the most similar, with 233 shared DEGs representing a 52.7% overlap (Fig. 1B). Both methods also produced a similar directionality in the gene expression changes (DESeq2: 212 downregulated, 134 upregulated; ALDEx3: 228 downregulated, 72 upregulated) (Supplementary Fig. 5A). The E20 vs. E16 comparison produced the largest absolute overlap (512 shared genes, 42.9% overlap), though DESeq2 still identified 630 exclusive DEGs while ALDEx3 identified 52 exclusive genes (Fig. 1C). In this case, there was less agreement between the methods regarding the directionality of gene expression with roughly equal numbers of up- and down-regulated genes for DESeq2 (595 downregulated, 547 upregulated) and a general down-regulation observed with ALDEx3 (121 upregulated, 443 downregulated) (Supplementary Fig. 5A).

**Figure 1:**
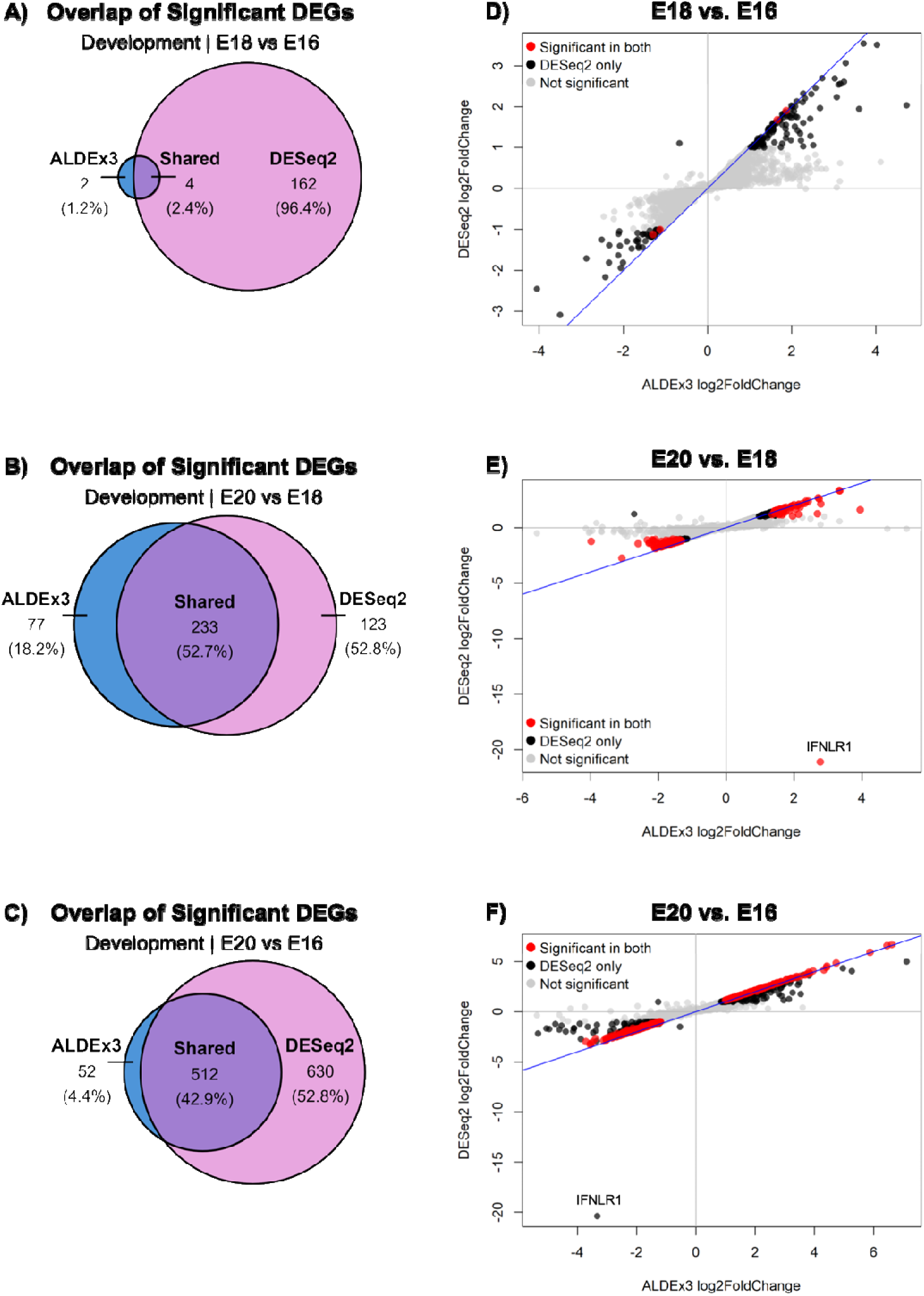
Differences between ALDEx3 and DESeq2 during development. Comparison of DEGs across three developmental comparisons: E18 vs. E16 (A, D), E20 vs. E18 (B, E), and E20 vs. E16 (C, F). **(A-C)** Venn diagrams showing the number and percentage of significant DEGs (log2 fold change ≥ 1, adj. p-value ≤ 0.05) identifie uniquely by ALDEx3, uniquely by DESeq2, or shared between both methods. **(D-F)** Scatter plots comparing log fold change values calculated by ALDEx3 (x-axis) and DESeq2 (y-axis), with blue line representing unity (x=y).

We analyzed the log2 fold-change values during development between methods to identify the consistency of individual gene expression, particularly for the DEGs that were only significant using DESeq2 and may have lower replicability. We found that, while DESeq2 identified more significant genes, the fold change estimates of the DEGs unique to DESeq2 were lower (i.e., more conservative) than the estimates made by ALDEx3 (Fig. 1D-F). However, the DEGs that were significant in both methods had generally similar fold change values. A small number of significant DEGs showed opposing fold-change signs between methods: *Them4* in the E18 vs. E16 comparison, *Ifnlr1* and *H2ac1* in the E20 vs. E18 comparison, and *Olr1* in the E20 vs. E16 comparison (Supplementary Table 1). Notably, Interferon Lambda Receptor 1 (*Ifnlr1*) was a consistent outlier in both the E20 vs. E18 and E20 vs. E16 comparisons, showing a sign change only in the E20 vs. E18 window (Fig. 1E-F).

Comparison of significant DEGs between rigid (DMB) or flaccid (PB) paralysis and the control samples showed a method-dependent difference in the total number of significant DEGs similar to the differences observed during development. With DMB compared to control, ALDEx3 identified only 3 unique DEGs (0.46%), with 201 shared between methods (30.97%) and 445 being exclusive to DESeq2 (68.57%; Fig. 2A). Both methods also described a transcriptomic response leaning toward downregulation (DESeq2: 404 downregulated, 242 upregulated; ALDEx3: 162 downregulated, 42 upregulated) (Supplementary Fig. 5B). For the PB comparison with controls, ALDEx3 identified no unique DEGs with all 3 genes in its dataset already present in the DESeq2 results (91 upregulated, 110 downregulated) (Fig. 2C, Supplementary Fig. 5B).

**Figure 2:**
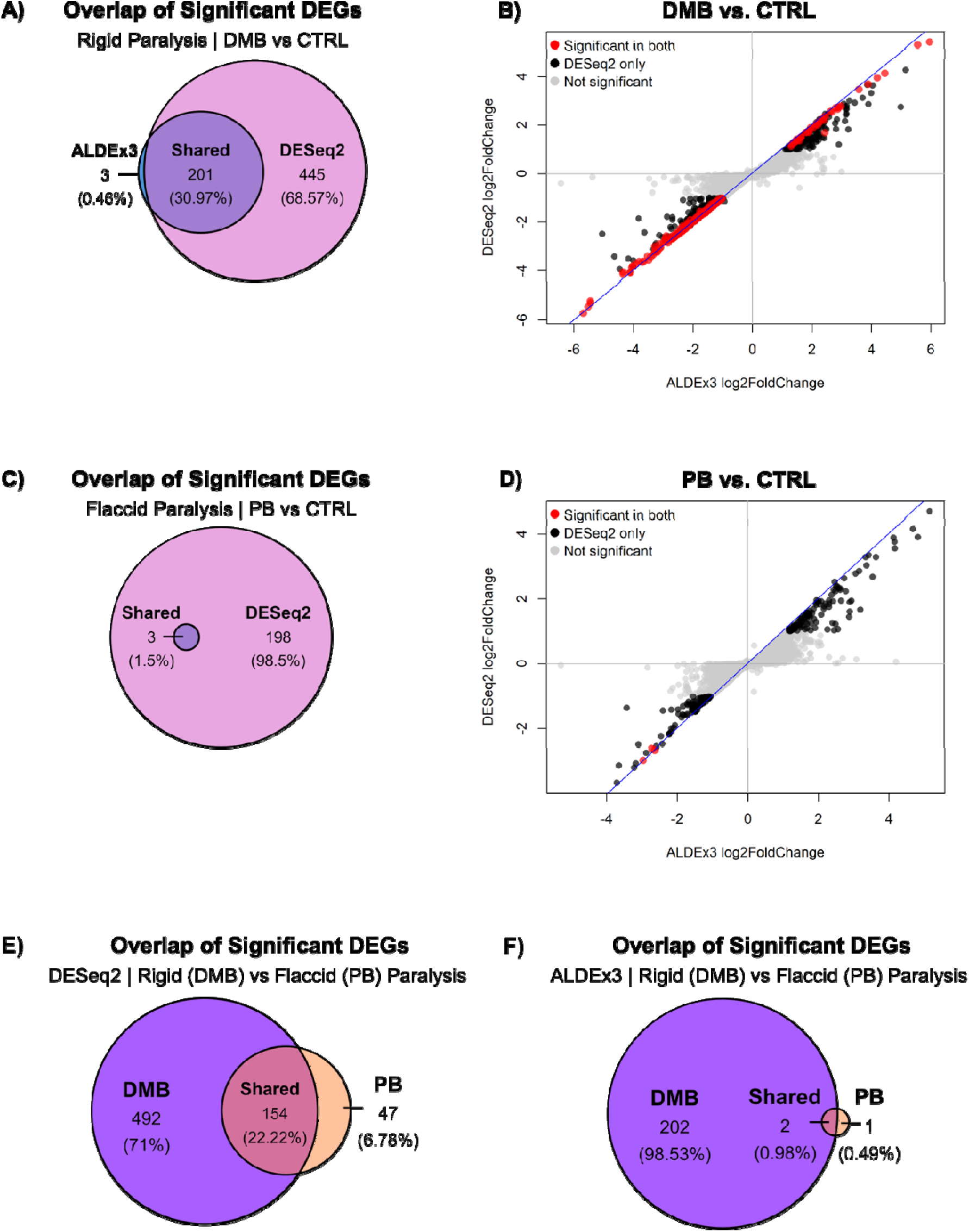
Differences between ALDEx3 and DESeq2 during paralysis. Comparison of DEGs across paralysis conditions: DMB vs. CTRL (A, B), PB vs. CTRL (C, D), and DMB vs. PB (E, F). **(A, C)** Venn diagrams showing the number and percentage of significant DEGs (log2 fold change ≥ 1, adj. p-value ≤ 0.05) identified uniquely by ALDEx3, uniquely by DESeq2, or shared between both methods for DMB and PB groups. **(B, D)** Scatter plots comparing log2 fold change values calculated by ALDEx3 (x-axis) and DESeq2 (y-axis) for the same two comparisons, with the blue line representing unity (x=y). **(E, F)** Venn diagrams showing the overlap of significant DEGs between DMB and PB groups, identified separately by DESeq2 (E) and ALDEx3 (F).

Analysis of the log2-fold change expression between methods also showed similar behavior to normal development, with both methods producing consistent fold changes for shared DEGs (Fig. 2B and D). Contrary to developmental comparisons, no significant DEGs showed opposite directionality in either paralysis comparison. As with development, genes significant only with DESeq2 had lower fold-change estimates than with ALDEx3, reinforcing that DESeq2 applies a more permissive significance threshold while accounting for scale uncertainty with ALDEx3 produces greater sensitivity (i.e., higher fold-change magnitudes)^40,41^. This was further supported by a sensitivity analysis of adjusted p-value thresholds for both ALDEx3 and DESeq2 (Supplementary Fig. 2), which showed that a substantial number of ALDEx3 genes became significantly differentially expressed at less restrictive threshold (adj. p < 0.10), while DESeq2 results were unaffected by this threshold change in any comparison.

When comparing the transcriptomic response between paralysis modes, DESeq2 identified 492 DEGs exclusive to DMB (71%), 47 exclusive to PB (6.78%), and 154 shared between paralysis modes (22.22%) (Fig. 2E). ALDEx3 identified a combined total of 205 DEGs, with DMB accounting for 98.53% of significant genes and minimal overlap with PB (2 shared, 0.98%) (Fig. 2F). Across both methods, DMB consistently produced a greater transcriptional response than PB, suggesting that rigid paralysis creates a broader transcriptional effect when compared to flaccid paralysis.

### 3.2 Shifts in Mechanosensing, ECM Remodeling, and Cytoskeletal Organization Observed with Late Embryonic Tendon Development

Between E16 and E18, Gene Set Enrichment Analysis (GSEA) of gene ontology biological processes (GOBP) for both ALDEx3 and DESeq2 datasets revealed upregulation of terms related to contractility, calcium ion transport, negative regulation of response to external stimulus, cytokine-mediated signaling, and cell-cell adhesion, while terms related to cellular response to stress and cell cycling were downregulated (Fig. 3). Based on the ALDEx3 results alone, we observed upregulation of terms related to mechanosensing (e.g., response to muscle stretch and detection of mechanical stimulus); actin-mediated cell contraction, movement, and organization; circadian rhythm; and calcium-mediated signaling (Fig. 3). GSEA of the ALDEx3 dataset did not identify any unique downregulated GOBP terms. GSEA of the DESeq2 dataset uniquely identified upregulated terms related to mechanosensing (e.g., response to laminar fluid shear stress, response to mechanical stimulus) and cell adhesion (e.g., cadherin-mediated cell-cell adhesion, heterotypic cell-cell adhesion, regulation of cell adhesion), as well as downregulated terms related to epigenetic gene expression and microtubule-based processes (Fig. 3). Collectively, these data suggest an active increase in mechanosensing and remodeling events between E16 and E18.

**Figure 3:**
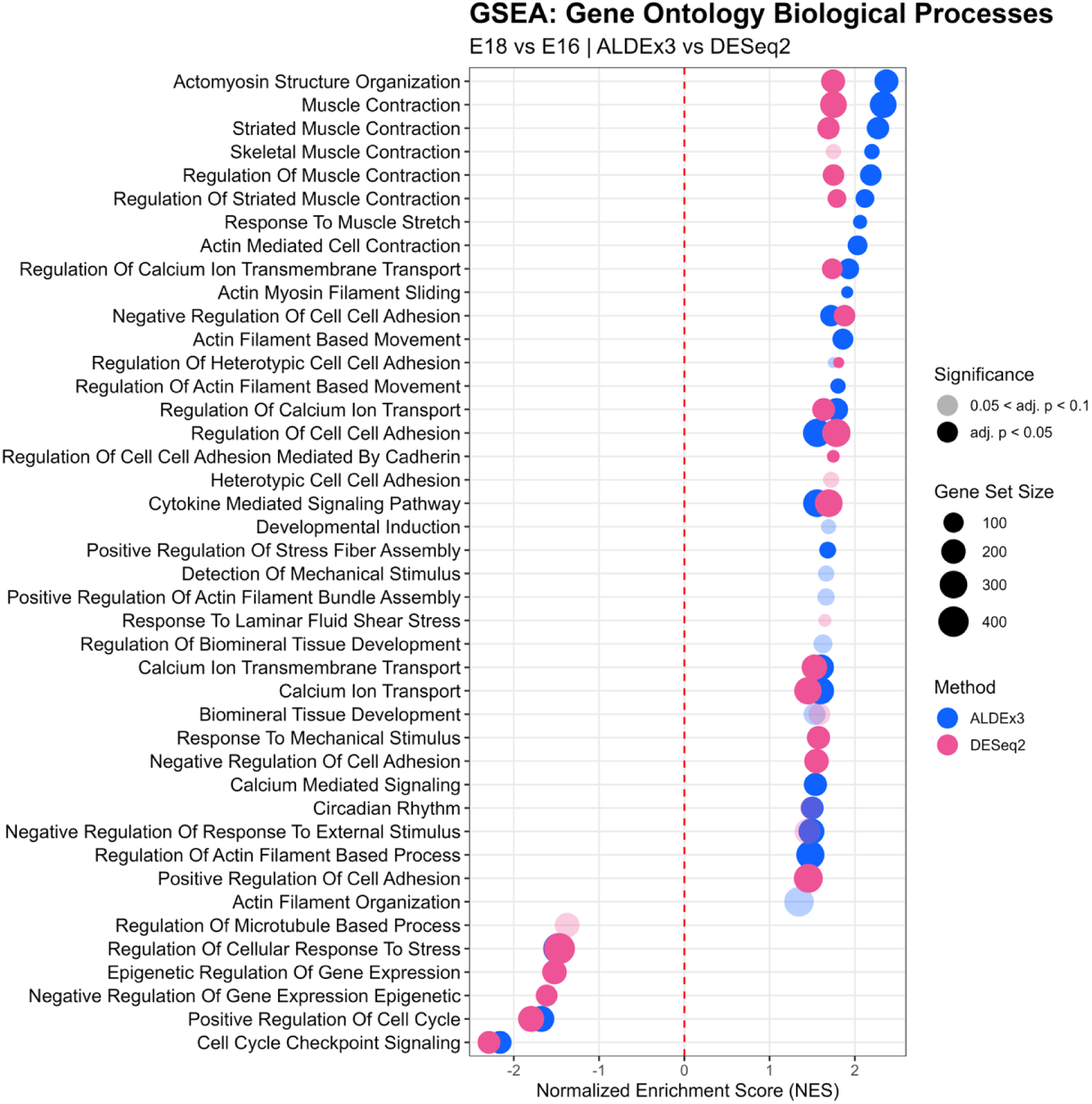
Gene set enrichment analysis of gene ontology biological process (GOBP) terms in the E18 vs. E16 comparison. Dot plot showing GSEA results for GOBP terms comparing ALDEx3 and DESeq2 for E18 vs. E16. Gene sets are ranked by normalized enrichment score (NES), with positive NES indicating enrichment at E18 and negative NES indicating enrichment at E16. The red dashed line indicates NES = 0.

These findings were further supported by GSEA of signaling pathways, which identified positive enrichment of muscle contraction, focal adhesion, circadian rhythm, Tgf-β receptor signaling in skeletal dysplasia, Stat3, Jak/Stat, and estrogen signaling pathways in both ALDEx3 and DESeq2 datasets, along with downregulation of keratan sulfate degradation, Hedgehog signaling, Rho GTPase-related signaling, gap junction assembly and trafficking, and cell cycle pathways (Fig. 4). Additionally, GSEA of signaling pathways in the ALDEx3 dataset alone identified upregulation of prostaglandin and Tgf-β receptor signaling, along with integrin-mediated cell adhesion, and downregulation of epigenetic regulation of gene expression (Fig. 4). Meanwhile, the DESeq2 dataset identified upregulation of extranuclear estrogen signaling, prostaglandin synthesis and regulation, exercise-induced circadian regulation, negative regulation of the Pi3k/Akt network, ECM organization, integrin-mediated cell adhesion, and regulation of Igf transport signaling, along with downregulation of Hedgehog “On State” signaling (Fig. 4). These results were consistent with the upregulation of mechanosensitive transcription factors Egr1 and Egr2 as well as the tendon-specific factor Mohawk, as determined from separate transcription factor enrichment analysis (Supplementary Fig. 6).

**Figure 4:**
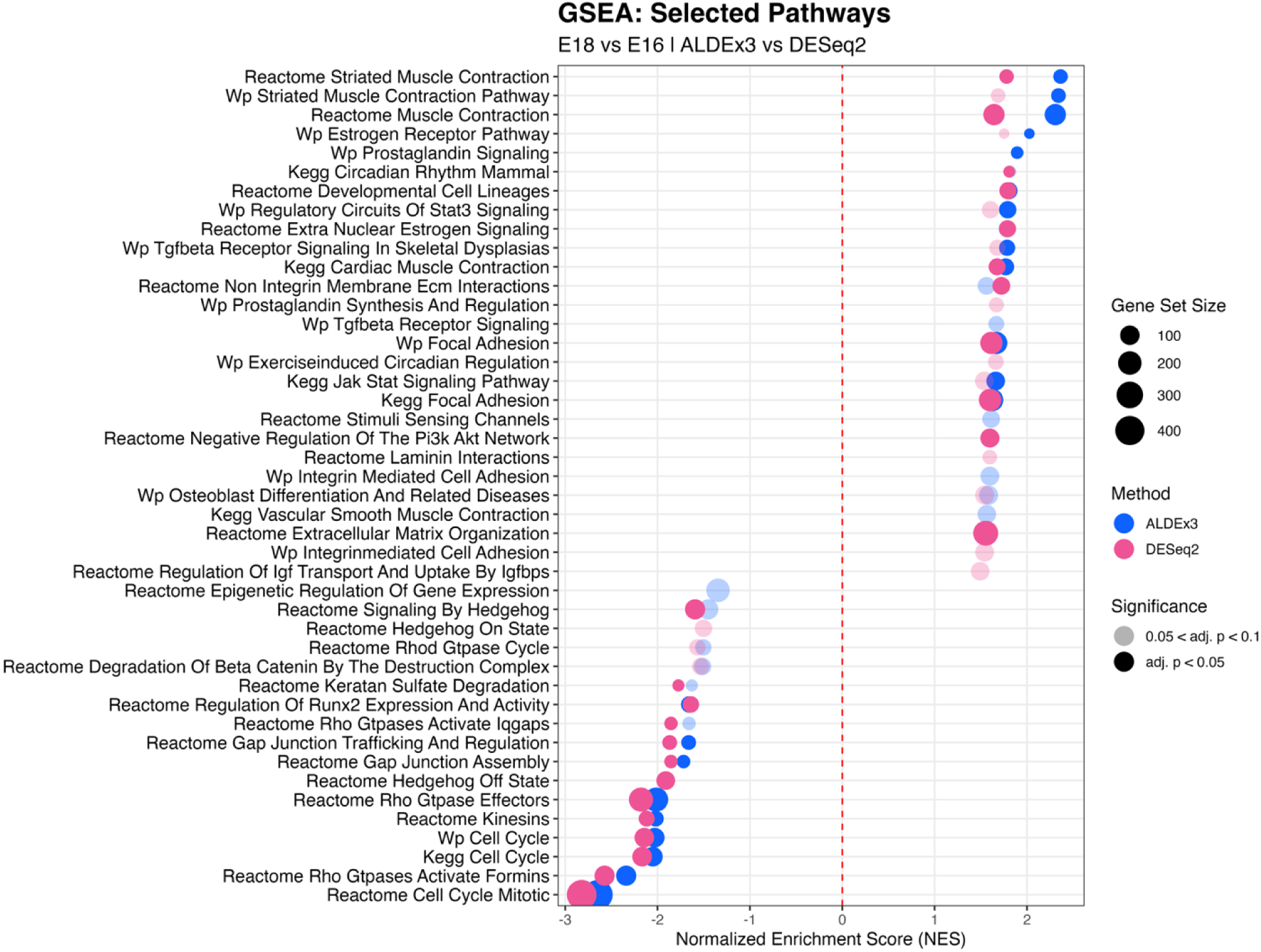
Gene set enrichment analysis of signaling pathways in the E18 vs. E16 comparison. Dot plot showing GSEA results for signaling pathways comparing ALDEx3 and DESeq2 for E18 vs. E16. Gene sets are ranked by normalized enrichment score (NES), with positive NES indicating enrichment at E18 and negative NES indicating enrichment at E16. The red dashed line indicates NES = 0.

During the E18 to E20 transition, GSEA of GOBPs for both the ALDEx3 and DESeq2 data revealed positive enrichment of negative regulation of response to external stimulus, regulation of stress fiber assembly, regulation of cell-cell and cell-substrate adhesion, and cytokine-mediated signaling pathways (Fig. 5), while microtubule processes, cytoskeleton organization, and cell cycling related terms were downregulated (Fig. 5). Downregulation of contractility-related processes and actin filament-based movement were also found uniquely with ALDEx3 (Fig. 5). Based on the DESeq2 data alone, we found upregulation of several ECM remodeling and matrix organization terms (e.g., ECM organization, connective tissue development, glycosaminoglycan biosynthetic and metabolic processes, collagen metabolism, tissue remodeling) (Fig. 5). We also observed upregulation of processes related to cell adhesion and mechanotransduction signaling (e.g., integrin-mediated signaling, cell-matrix adhesion, Jak-Stat receptor signaling, response to external stimulus, and the MAPK cascade terms P38Mapk, Jnk, and Erk1/Erk2) (Fig. 5). The DESeq2 data also indicated downregulation of microtubule-based movement and polymerization, protein localization to cytoskeleton, and negative regulation of microtubule depolymerization.

**Figure 5:**
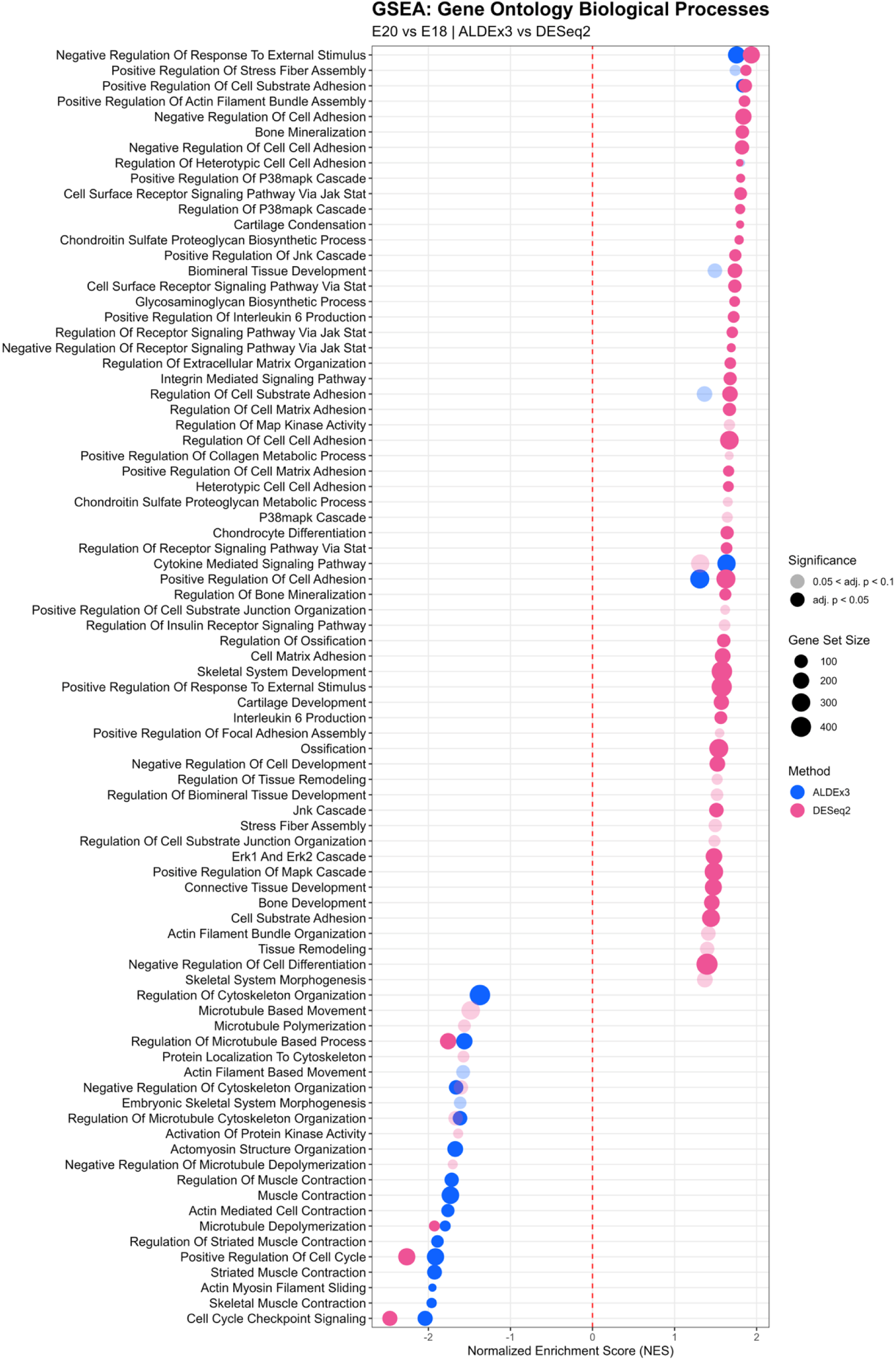
Gene set enrichment analysis of gene ontology biological process (GOBP) terms in the E20 vs. E18 comparison. Dot plot showing GSEA results for GOBP terms comparing ALDEx3 and DESeq2 for E20 vs. E18. Gene sets are ranked by normalized enrichment score (NES), with positive NES indicating enrichment at E20 and negative NES indicating enrichment at E18. The red dashed line indicates NES = 0.

These findings were further supported by GSEA of signaling pathways, which showed positive enrichment of Il-6 family signaling and downregulation of cell cycle and Rho GTPase-related pathways across both ALDEx3 and DESeq2 datasets (Fig. 6). ALDEx3 uniquely found positive enrichment of signaling by interleukins (e.g., Il-6) and Jak-Stat signaling (Fig. 6), whereas muscle contraction, Tgf-β receptor signaling, kinesins, and Smad2/3/Smad4 transcriptional activity were downregulated (Fig. 6). The DESeq2 data uniquely indicated positive enrichment of Mapk, focal adhesion through Pi3k/Akt/mTOR, Igf, Hippo, and Tgf-β receptor signaling. The DESeq2 data also found negative enrichment of STAT3 and other interleukin signaling pathways (Fig. 6). Finally, upregulation of Mohawk was identified with transcription factor overrepresentation analysis (Supplementary Fig. 7).

**Figure 6:**
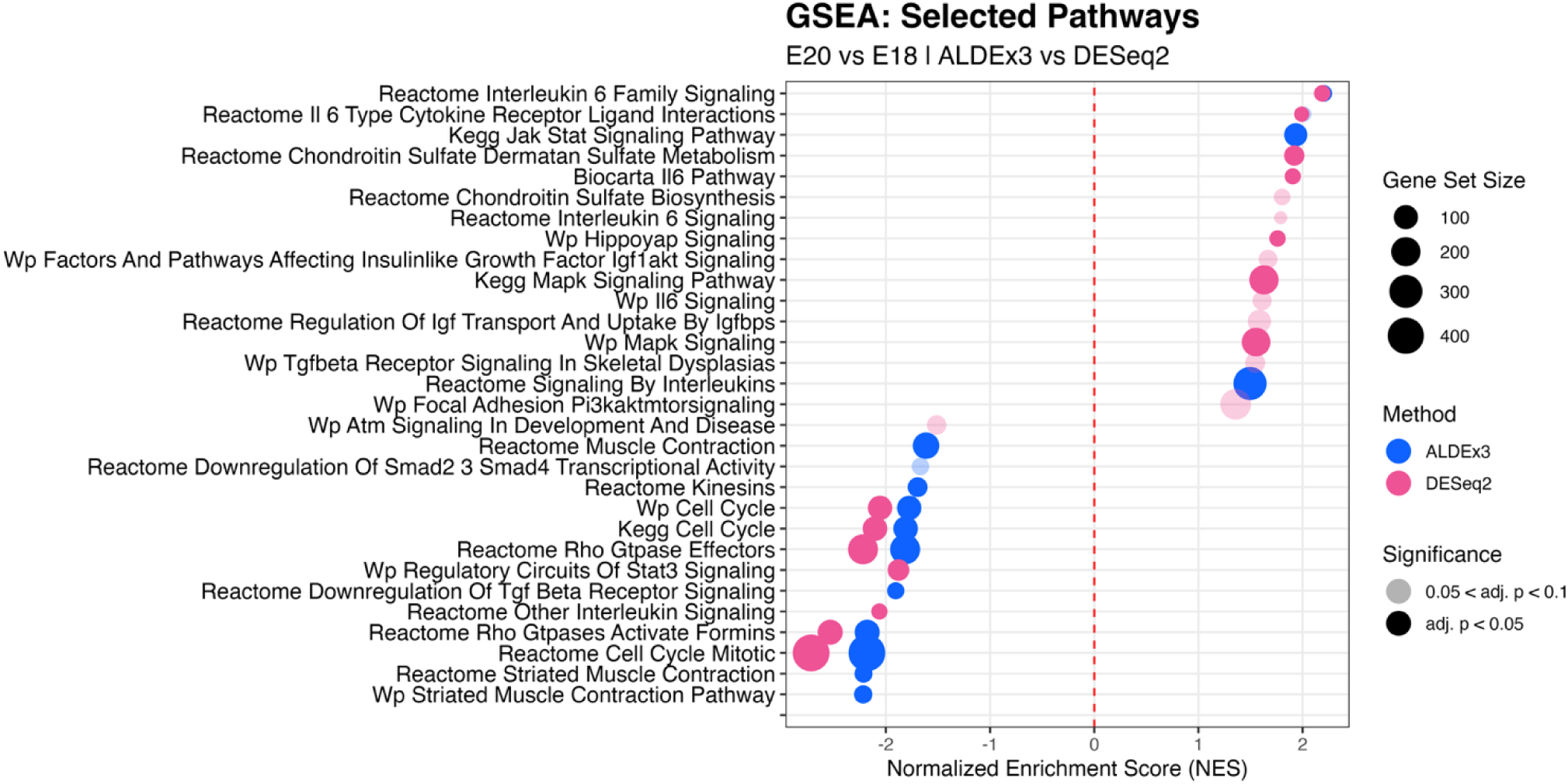
Gene set enrichment analysis of signaling pathways in the E20 vs. E18 comparison. Dot plot showing GSEA results for signaling pathways comparing ALDEx3 and DESeq2 for E20 vs. E18. Gene sets are ranked by normalized enrichment score (NES), with positive NES indicating enrichment at E20 and negative NES indicating enrichment at E18. The red dashed line indicates NES = 0.

Looking holistically across the complete E16 to E20 transition, GSEA of GOBPs revealed positive enrichment of mechanosensing and physical stimulus response terms (e.g., response to laminar fluid shear stress, response to mechanical stimulus, response to external stimulus), Jak/Stat signaling, as well as cell adhesion-related terms (e.g., cell surface receptor signaling via Stat, cell-substrate adhesion, actomyosin structure organization), and negative enrichment of epigenetic regulation of gene expression terms, microtubule-based processes (e.g., microtubule depolymerization, microtubule cytoskeleton organization), and cell cycling-related terms, across both DESeq2 and ALDEx3 datasets (Fig. 7). The ALDEx3 dataset uniquely identified upregulation of calcium ion transport-related terms and downregulation of embryonic skeletal system morphogenesis (Fig. 7). Based on the DESeq2 data alone, we observed upregulation of terms related to ECM organization and adhesion (e.g., ECM organization, ECM assembly, cell-substrate adhesion, integrin-mediated signaling, positive regulation of cell-cell adhesion), as well as terms related to stimulus and stress response signaling (e.g., integrated stress response signaling, detection of stimulus, response to Tgf-β). Additional upregulated terms in the DESeq2 data included entrainment of circadian clock, p38 Mapk signaling, Mapk activity, circadian regulation of gene expression and rhythm, and tissue homeostasis, while downregulated terms included microtubule-based movement and negative regulation of cytoskeleton organization (Fig. 7).

**Figure 7:**
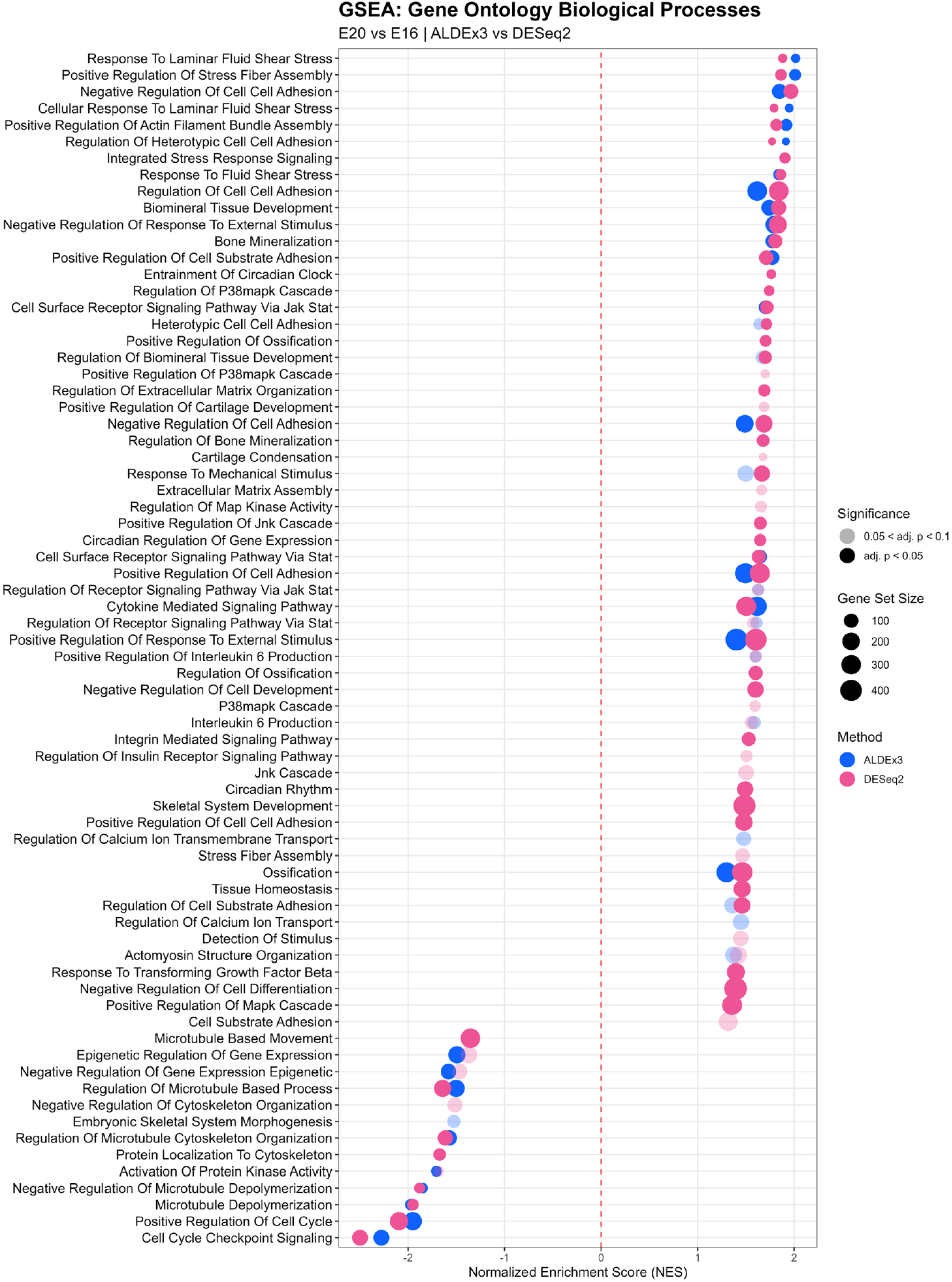
Gene set enrichment analysis of gene ontology biological process (GOBP) terms in the E20 vs. E16 comparison. Dot plot showing GSEA results for GOBP terms comparing ALDEx3 and DESeq2 for E20 vs. E16. Gene sets are ranked by normalized enrichment score (NES), with positive NES indicating enrichment at E20 and negative NES indicating enrichment at E16. The red dashed line indicates NES = 0.

These findings were further supported by GSEA of signaling pathways, which identified positive enrichment of Il-6 family signaling, nuclear and estrogen receptor pathways, extranuclear estrogen signaling, non-integrin membrane-ECM interactions, focal adhesion, and Igf transport in both ALDEx3 and DESeq2 results, as well as downregulation of Rho GTPase-related terms, kinesins, and cell cycling-related terms (Fig. 8). ALDEx3 uniquely identified positive enrichment of Il-6 cytokine receptor interactions. Paradoxically, ALDEx3 uniquely identified a downregulation of “downregulation of Tgf-β receptor signaling” (i.e., an upregulation) and downregulation of canonical and non-canonical Tgf-β signaling (Fig. 8), suggesting a complicated role of Tgf-β during development. Notably, the DESeq2 dataset identified upregulation of Tgf-β receptor signaling (Fig. 8). Additionally, DESeq2 results included upregulation of terms related to ECM and connective tissue signaling (e.g., ECM-receptor interaction, chondroitin sulfate biosynthesis and metabolism, ECM organization), as well as Hippo, Il-6 signaling, circadian rhythm and clock-related pathways, focal adhesion (both independently and through Pi3k/Akt/mTOR signaling), and muscle contraction, and did not uniquely identify downregulation of any pathway (Fig. 8). These results were supported by upregulation of Smad1, Smad3, Mohawk, Tead1, and Tead3 identified in the transcription factor enrichment analysis results (Supplementary Fig. 8).

**Figure 8:**
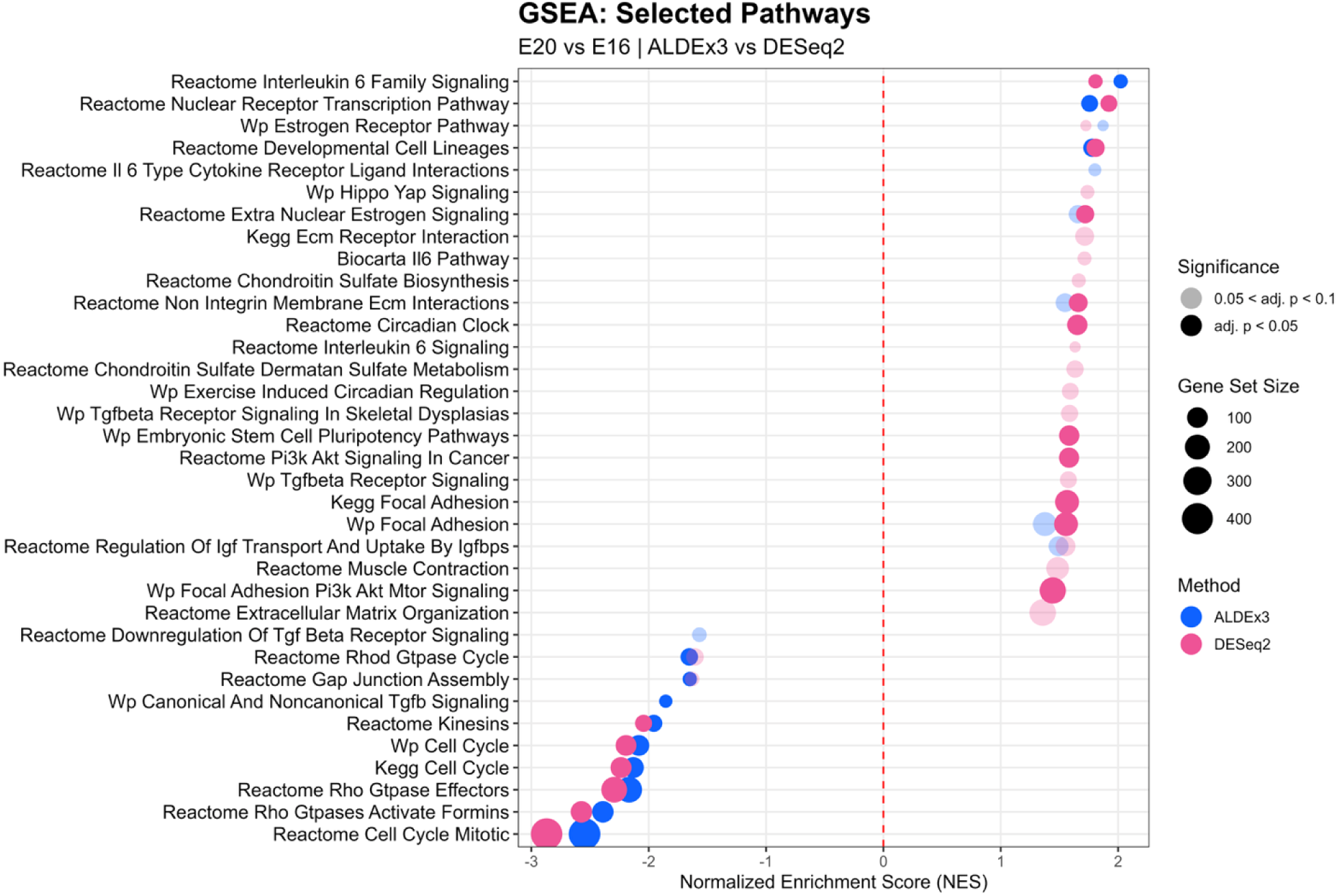
Gene set enrichment analysis of signaling pathways in the E20 vs. E16 comparison. Dot plot showing GSEA results for signaling pathways comparing ALDEx3 and DESeq2 for E20 vs. E16. Gene sets are ranked by normalized enrichment score (NES), with positive NES indicating enrichment at E20 and negative NES indicating enrichment at E16. The red dashed line indicates NES = 0.

### 3.3 Rigid and Flaccid Paralysis Produce Mode-Dependent Transcriptomic Responses

#### Characterized by Impaired Mechanosensing, ECM Remodeling, and Contractility

GSEA of GOBPs revealed that the transcriptomic response to paralysis was fundamentally dependent on paralysis mode, given that most terms in the rigid paralysis (DMB) comparison were negatively enriched, in contrast to the predominantly positive enrichment observed in flaccid paralysis (PB) (Fig. 9). Signal release was the only positively enriched term shared between both paralysis modes, identified by ALDEx3 in PB and by DESeq2 in DMB. In rigid paralysis, both ALDEx3 and DESeq2 identified downregulation of terms associated with cytoskeleton and actin filament organization, stem cell development and differentiation, tissue morphogenesis, microtubule dynamics, and proteoglycan biosynthetic processes, while ALDEx3 alone additionally identified downregulation of calcium ion transport, stem cell differentiation, regulation of cell cycle processes, and actomyosin structure organization (Fig. 9). DESeq2 did not uniquely identify any additional GOBP terms in the rigid paralysis comparison.

**Figure 9:**
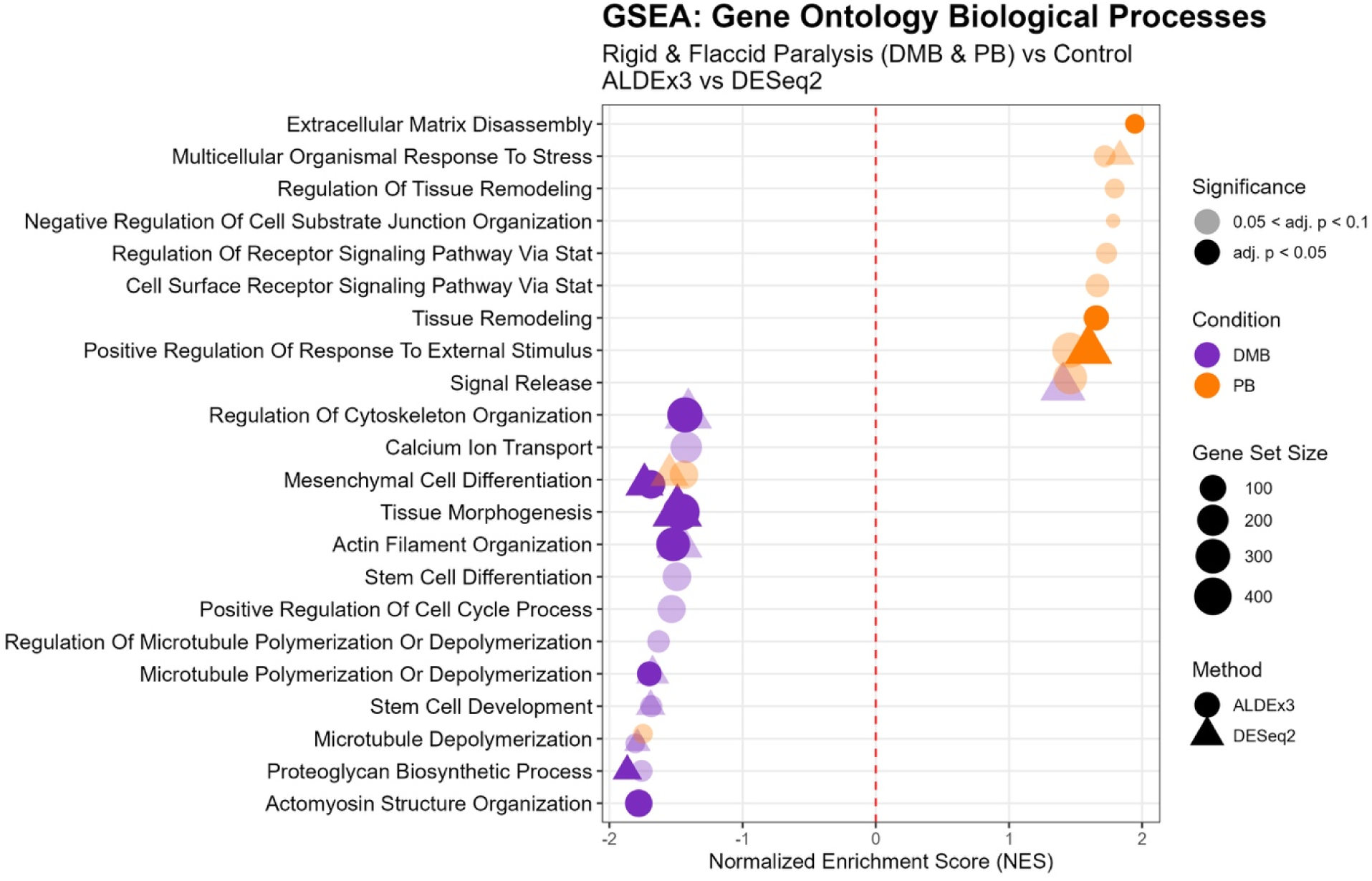
Gene set enrichment analysis of gene ontology biological process (GOBP) terms in the DMB and PB vs. CTRL comparisons. Dot plots showing GSEA results for GOBP terms comparing ALDEx3 and DESeq2 for rigid (DMB) and flaccid (PB) paralysis, each relative to control. Gene sets are ranked by normalized enrichment score (NES), with positive NES indicating enrichment in the paralysis condition and negative NES indicating enrichment in control. The red dashed line indicates NES = 0.

These findings were further supported by signaling pathway enrichment analysis, which identified upregulation of epigenetic regulation of gene expression and Jak-Stat signaling in both paralysis modes across both datasets, while Wnt ligand biogenesis and trafficking and Hedgehog signaling were downregulated in both paralysis modes across both datasets (Fig. 10). In rigid paralysis specifically, both DESeq2 and ALDEx3 identified broad downregulation of ECM remodeling pathways, including significant negative enrichment of collagen formation and biosynthesis, ECM organization, and glycosaminoglycan (GAG) metabolism, along with downregulation of the Wnt signaling pathway and Rho GTPase-related terms (Fig. 10); the Wnt downregulation was consistent with reduced activity of the Wnt transcriptional regulators Lef1 and Tcf4 identified by transcription factor analysis (Supplementary Fig. 9), while the ECM findings were supported by downregulation of Runx2 and Smad3 from the same analysis (Supplementary Fig. 9). Additionally, ALDEx3 uniquely identified downregulation of contractility-related pathways in rigid paralysis, including smooth muscle contraction and striated muscle contraction (Fig. 10). DESeq2 uniquely identified downregulation of keratan sulfate metabolism, elastic fiber formation, and GAG biosynthesis in rigid paralysis (Fig. 10). Notably, Mohawk showed activity with DMB treatment in both the upregulated and downregulated gene sets identified by transcription factor analysis (Supplementary Fig. 9), suggesting complex dysregulation of tendon-specific transcriptional activity with rigid paralysis. Collectively, these data suggest a comprehensive impairment of mechanotransduction and structural ECM integrity in response to rigid paralysis.

**Figure 10:**
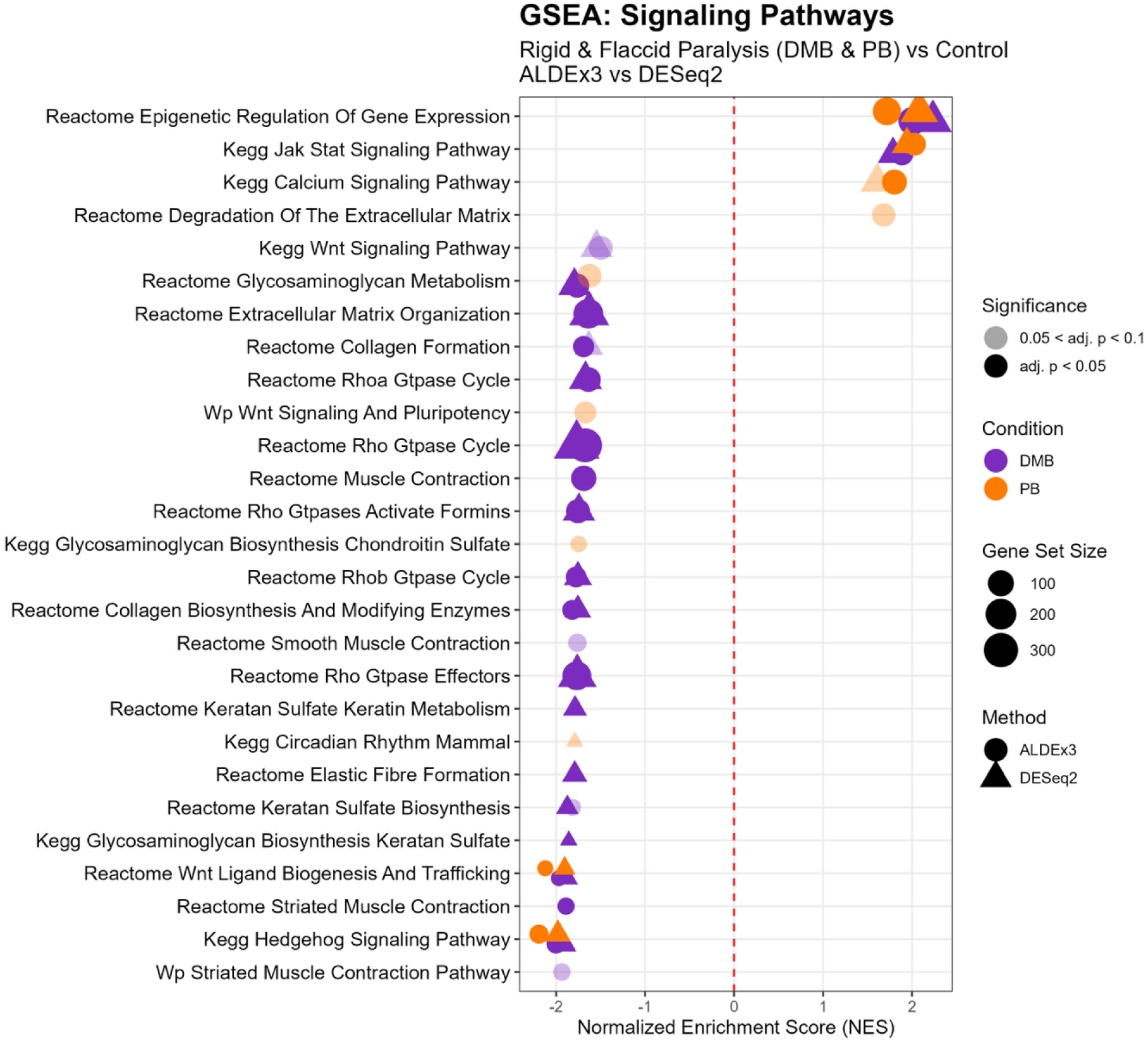
Gene set enrichment analysis of signaling pathways in the DMB and PB vs. CTRL comparisons. Dot plots showing GSEA results for signaling pathways comparing ALDEx3 and DESeq2 for rigid (DMB) and flaccid (PB) paralysis, each relative to control. Gene sets are ranked by normalized enrichment score (NES), with positive NES indicating enrichment in the paralysis condition and negative NES indicating enrichment in control. The red dashed line indicates NES = 0.

The flaccid paralysis (PB) comparison revealed positive enrichment of GOBP stress response terms in both ALDEx3 and DESeq2 datasets, including multicellular organismal response to stress and positive regulation of response to external stimulus (Fig. 9). ALDEx3 additionally identified unique upregulation of ECM disassembly, tissue remodeling, cell-substrate junction organization, and cell surface receptor signaling via Stat, as well as downregulation of microtubule depolymerization, consistent with the downregulation of microtubule-based processes observed across developmental transitions (Fig. 9). DESeq2 did not identify any unique GOBP terms in the PB comparison.

These findings were further supported by GSEA of signaling pathways, which identified positive enrichment of calcium signaling in both datasets, in addition to the epigenetic regulation and Jak-Stat signaling upregulation noted above across both paralysis modes (Fig. 10); this was supported by upregulation of Stat5a and Stat3 from transcription factor analysis (Supplementary Fig. 10). Wnt signaling and its ligand biogenesis and trafficking were negatively enriched in both datasets (Fig. 10), consistent with the downregulation observed in rigid paralysis and with reduced activity of the Wnt transcriptional regulators Lef1 and Tcf7 identified by transcription factor enrichment analysis (Supplementary Fig. 10); however, concurrent upregulation of Tcf7l2 suggests more complex regulation of Wnt transcriptional activity in flaccid paralysis. ALDEx3 uniquely identified downregulation of GAG-related terms, including GAG metabolism and GAG biosynthesis of chondroitin sulfate, as well as downregulation of Wnt signaling and pluripotency pathways (Fig. 10). DESeq2 uniquely identified downregulation of circadian rhythm in mammals, contrasting with the upregulation observed in developmental comparisons (Fig. 10).

## 4. Discussion

In this study, we used two analytical techniques to identify the transcriptomic changes and biological processes associated with late tendon maturation in the chick embryo model. The results from both analyses were generally consistent and suggest that the largest shift in differential gene expression occurs between E16 and E20 and that, among the paralysis conditions, rigid paralysis (DMB) produces the greatest effect relative to vehicle controls. Surprisingly, the fewest number of DEGs were found between E16 and E18, which is when the largest gains in tendon mechanics occur^23,25^. Additionally, despite the different transcriptomic response due to DMB and PB treatment, our previous data found that both rigid and flaccid paralysis produce similar decreases in tendon mechanical properties^23,25^. These findings suggest that there are numerous biological pathways being regulated that do not directly influence changes in tendon mechanics. Focusing on the smaller set of DEGs between E16 and E18, we find that upregulation of circadian rhythm, Tgf-β signaling, focal adhesion signaling, contractility, and response to mechanical stimulus and ECM organization overlap with the largest increase in mechanical properties. When looking holistically across the earliest and latest timepoints in our data (E20 vs. E16), Hippo, circadian clock, Tgf-β, and focal adhesion signaling are all positively regulated with development and may be playing a broader role in tendon development. Across the paralysis conditions, Jak/Stat upregulation and Wnt signaling downregulation are shared between rigid and flaccid paralysis. As these candidate pathways will require functional validation, future work can perturb them to identify their unique importance in driving late tendon development and test whether exogeneous manipulation in tendon constructs can improve tendon tissue engineering approaches.

Our findings both agree and disagree with prior studies of tendon development. Changes in Tgf-β signaling and pathways related to ECM organization were also reported in chicks during early embryonic tendon formation (i.e., E0 to E7 and E11-17)^32,36^; however, the upregulation in muscle contraction we observed throughout late development is opposite to what was seen previously in E11 - E17 chicken tendons^36^. Our data also resembles some of the findings seen in early zebrafish tendon development^57^. In mouse tendons, circadian rhythm has also previously been shown to regulate collagen homeostasis and the tenocyte transcriptomic response^58–60^, supporting a conserved pattern for this pathway across species. Notably, our results are inconsistent with findings in postnatal mouse tendon, where Hippo, Wnt, and Tgf-β signaling, along with ECM organization, are downregulated over comparable developmental stages^37^. This could potentially be due to differences in loading between species during this time, where chicks experience isometric loading due to confinement of the eggshell while mice are able to freely ambulate post-natally^61,62^. Additionally, we found that focal adhesion signaling increased with development across all developmental comparisons, with focal adhesion Pi3k/Akt/mTOR signaling upregulated in the E20 vs. E18 and E20 vs. E16 comparisons. Fak sits upstream of Pi3k/Akt^63^, and Pi3k/Akt signaling has been shown to act as a mechanotransducer influencing tenogenic differentiation of tendon-derived stem cells in response to uniaxial mechanical stimulation^64^ and neonatal tendon regeneration^65^. Our data are consistent with a prior study of transgenic mice showing that Fak knockout alters tendon size and cell composition; however, Fak knockout did not decrease the tendon material properties, suggesting that Fak is important for growth but not tissue construction^38^. It is also notable that terms associated with cell cycling and microtubule organization were downregulated at later developmental timepoints, which is consistent with decreased proliferation and an overall reduction in cell volume fraction^17,18^. Collectively, our data add to the available literature regarding the biological signaling that changes during tendon development and highlight that some pathways are conserved and others diverge across timepoints and species.

A unique feature of our study is that we used two statistical models to determine differential gene expression. Specifically, we utilized ALDEx3 in addition to DESeq2 to model the uncertainty resulting from the unobserved total gene expression (i.e., differences in means between timepoints or conditions), which has been shown to reduce the number of false positives and, in some circumstances, change the estimated sign of the log fold change^40,41^. For both the developmental and paralysis datasets, we generally found consistent findings between both analytical models. However, we also observed distinct patterns in the directionality of the transcriptional response at each developmental stage. In particular, the E20 vs. E16 comparison showed the greatest discrepancy between methods, where ALDEx3 suggested a downregulation in gene expression, while DESeq2 showed roughly equal numbers of up- and downregulated genes. This discrepancy, along with instances of fold-change sign flips for individual genes (Supplementary Table 1), is likely due to the different ways DESeq2 and ALDEx3 account for the unknown differences in total gene expression across samples. Specifically, DESeq2 implicitly assumes that the change in total expression between conditions is approximately equal to the median fold-change across all genes whereas ALDEx3 assumes no difference in total expression between conditions and propagates uncertainty in this assumption through downstream inference^39,40,51^. In addition to broad differences in gene regulation, we found some unique differentially regulated pathways and genes from each analytical technique. Notably, Interferon Lambda Receptor 1 (Ifnlr1, also known as Il28rA) was found as a consistent outlier between methods in both the E20 vs. E18 and E20 vs. E16 comparisons. Ifnlr1 is a class II cytokine receptor that binds type III interferons (Ifn-λ) to activate Jak/Stat signaling^66–68^. Ifnlr1 has been reported to be broadly expressed across tissues, including skeletal muscle, and is also present in embryonic stem cells^67–69^, which could partially explain its presence in our dataset. To ensure that this outlier did not artificially inflate downstream Jak/Stat signaling enrichment, we repeated the analysis with Ifnlr1 excluded and found no change in the results. Therefore, the use of both DESeq2 and ALDEx3 highlights the subsets of our data that are robust and sensitive to scale uncertainty as well as to the analytical technique itself.

A main interest of this study was to identify potential biological mechanisms that may explain the mechanosensitivity of tendon development. At the GO term level, comparisons involving embryonic day 16 (E18 vs. E16 and E20 vs. E16) revealed that the mechanical stimulus response and contractility observed broadly across these windows reflects enrichment of specific biological processes including mechanotransduction, mechanical stimulus detection, and cell adhesion. Similarly, contractility-related pathways were also positively enriched between E16 and E18. These data suggest that tenocytes are responding to both intrinsic matrix tension and extrinsic mechanical cues during this period. Notably, many of the contractility terms were annotated as muscle-associated processes; however, upon closer examination of the specific gene sets driving this enrichment (for full gene list see Supplemental Data File 3), we interpret this signal as a combination of two sources. While a subset of the contributing genes is specific to muscle tissue, suggesting potential muscle contamination in our samples, many of the significant genes present in these gene sets are associated with generic cell contractility rather than muscle-specific function, which likely reflects the contractile activity of tenocytes. This contractile activity, alongside extrinsic tensile cues, has been shown to be essential for maintaining tendon tissue homeostasis and supporting embryonic development^70–73^. Interestingly, in the later E20 vs. E18 comparison, gene ontology results suggest a downregulation in contractility and regulation of contraction terms, which could be partially attributed to a reduction in unilateral and bilateral limb movements *in ovo* between embryonic day 18 and hatching due to confinement by the eggshell^61^. Rho GTPase–related pathways also appeared to be downregulated in all developmental comparisons, which is counterintuitive given that Rho GTPases regulate cell contractility^74^ and contractility-related biological processes were enriched during this same time period. This suggests that alternate mechanisms may be responsible for the possible increased cell contractility generally seen during development.

Our pathway analysis also identified a complex trend in Tgf-β signaling across development. Between E16 and E18, ALDEx3 suggested an upregulation of Tgf-β receptor signaling and a downregulation of this same pathway between E18 and E20. Interestingly, this was also observed in mice, with a transient surge in Tgf-β1 at P7, which then decreased at later developmental stages^37^. Additionally, in our overall E20 vs. E16 comparison, we found conflicting Tgf-β enrichment depending on the database used: GOBP analysis identified upregulation of response to Tgf-β in the DESeq2 dataset, while pathway analysis identified upregulation of Tgf-β receptor signaling. Notably, pathways analysis also showed significant downregulation of both canonical and non-canonical Tgf-β signaling overall in the ALDEx3 dataset. This may be due to differential activity of Tgf-β receptors since prior mouse data found intermittent differential expression of Tgf-βr1, whereas Tgf-βr3 was upregulated between P0 and P35^37^. Together, these discrepancies suggest that the role of Tgf-β in late tendon development is dynamic, complex, and worthy of further investigation.

Regarding Hippo signaling, we found that it was consistently upregulated in the E20 vs. E18 and E20 vs. E16 comparisons. This agrees with prior data of chick tendons at earlier embryonic stages (E11 – E17) and our own data covering E14 – E20^19^. Despite these findings, conditional knockout of *Yap1* at postnatal stages in mice only affected collagen expression and matrix organization without significantly altering tendon ultrastructural or mechanical properties^37^, suggesting that Hippo signaling through Yap1 regulation does not contribute to the mechanical changes observed in tendons during late development. However, further studies are necessary since the Yap homologue Taz was not knocked out in this model and may compensate for the loss of Yap. Similarly, previous studies have found elevated Yap signaling in tendon engineered constructs compared to E17 tendons^36^, indicating that construct maturation is not limited by insufficient Yap activity. Together, these findings suggest that while Yap may play a role in tendon development, it does not appear to be a critical driver of functional development or a limiting factor for improving tendon construct function.

Regarding our findings following musculoskeletal paralysis, Jak/Stat signaling was interestingly the only pathway consistently upregulated in both developmental and paralysis comparisons (Fig. 4 and 6). Consistent with the importance of Jak/Stat signaling during tendon development, treatment of engineered human ligaments with Jak1 increased collagen content and maximum tensile load^75^. Additionally, inhibition of Jak2 in juvenile rats stunted growth, postnatal development, and reduced the maximum tensile load of the Achilles and patellar tendons^76^. Furthermore, the upregulation of cellular stress response in our paralysis gene ontology enrichment analysis is consistent with the established role of Jak/Stat in mediating cellular responses to biological stress^77,78^.

More broadly, we found that the transcriptomic response of flaccid paralysis (PB) was smaller in magnitude than that of rigid paralysis (DMB). Interestingly, while DMB showed a greater overall number of enriched GOBP terms, the majority of these were downregulated, whereas PB showed a predominantly upregulated transcriptomic response, this is unexpected but could be due to the activation of abnormal cellular processes. For example, among the upregulated processes in PB were positive regulation of response to external stimulus and tissue remodeling, considering these alongside the upregulation of stress response terms, we interpret that this represents a cellular stress response to unloading. In contrast, rigid paralysis (DMB) produced downregulation of muscle contraction, Rho GTPase, ECM organization, and actomyosin structure organization terms. These differences in the transcriptomic response between rigid and flaccid paralysis likely reflect the tension produced in the tendons with paralysis, with rigid paralysis generating a static load and flaccid paralysis unloading the tissue. Despite these differences, Wnt signaling, which a known mechanotransduction pathway^79^, was downregulated under both paralysis conditions. Canonical Wnt signaling has been shown to suppress *Scx*, *Mkx*, and *Tnmd* expression, thus acting as a repressor of tenogenic gene expression^34,35^; however, recent data also suggests that Wnt plays a role in tenocyte development and morphogenesis at tendon attachments in zebrafish during early development^80^. Therefore, the consistent downregulation of Wnt across both paralysis conditions may reflect an attempt by the cells to compensate for mechanical dysregulation by promoting tenogenic gene expression; however, it is also possible that Wnt downregulation is a mechanism explaining impaired tendon development with paralysis^23,25^.

In conclusion, this study identified key candidate pathways potentially driving late tendon development. In particular, we found that Hippo, Pi3k/Akt/mTOR, Tgf-β, and focal adhesion signaling are mechanotransduction pathways that may be important for producing a robust tensile load-bearing tissue. Furthermore, we found that both rigid and flaccid paralysis treatments were associated with impaired mechanotransduction, impaired ECM remodeling and organization pathways, and an increased cellular stress response. These findings further support that mechanical loading plays a critical role in regulating late tendon development. In the future, additional perturbation and tissue function studies are necessary to confirm whether these pathways are truly critical for late tendon development. One limitation to this study is that the chicken genome is poorly annotated relative to the mouse and human genomes. To improve our downstream analysis (GSEA), we mapped the chicken genes to human orthologs, which reduced the total number of genes available for our gene enrichment analysis by approximately 25%. Therefore, additional studies investigating similar developmental timepoints in mice are important to further establish the pathways that are conserved across species. This would also strengthen the identification of which pathways may be most likely to improve tendon/ligament tissue engineering such that future tissue constructs can truly replicate tendon and ligament function.

## Acknowledgements

Funding for this project was provided by the National Institutes of Health (R21 AR075941) and the US-Ireland R&D Partnership (NSF 2127527) with partner funding from Research Ireland (20/US/3718) and from the Department for the Economy Northern Ireland (USI 180). We would like to thank the Penn State Poultry Education Research Center (PERC) for supplying the fertilized eggs for this study.

## Supplementary Figures

**Supplementary Figure 1.**
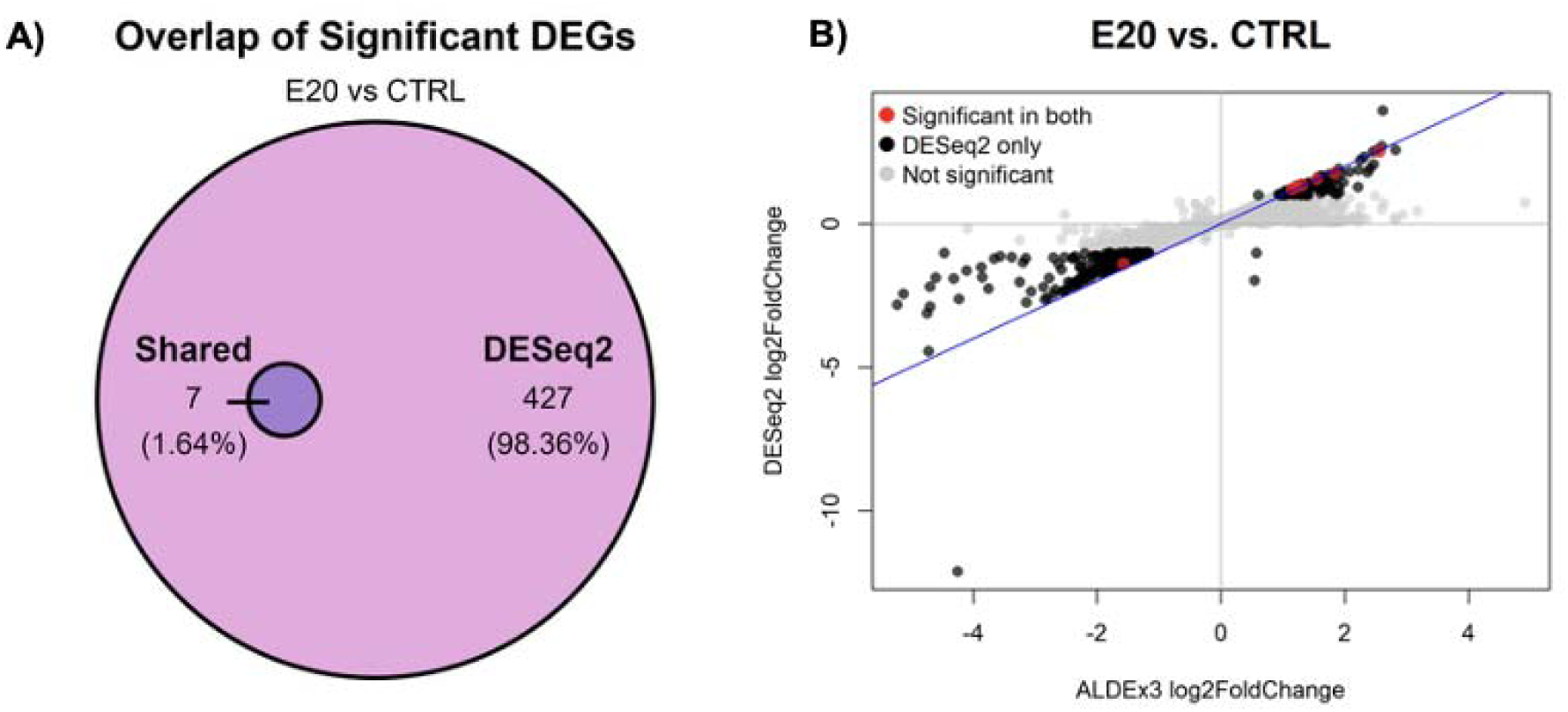
Differences between ALDEx3 and DESeq2, E20 vs. CTRL. **A)** Venn diagram showing the number and percentage of significant DEGs (log2 fold change ≥ 1, adj. p-value ≤ 0.05) identified uniquely by ALDEx3, uniquely by DESeq2, or shared between both methods. **B)** Scatter plot comparing log2 fold change values calculated by ALDEx3 (x-axis) and DESeq2 (y-axis), with the blue line representing unity (x=y).

**Supplementary Figure 2:**
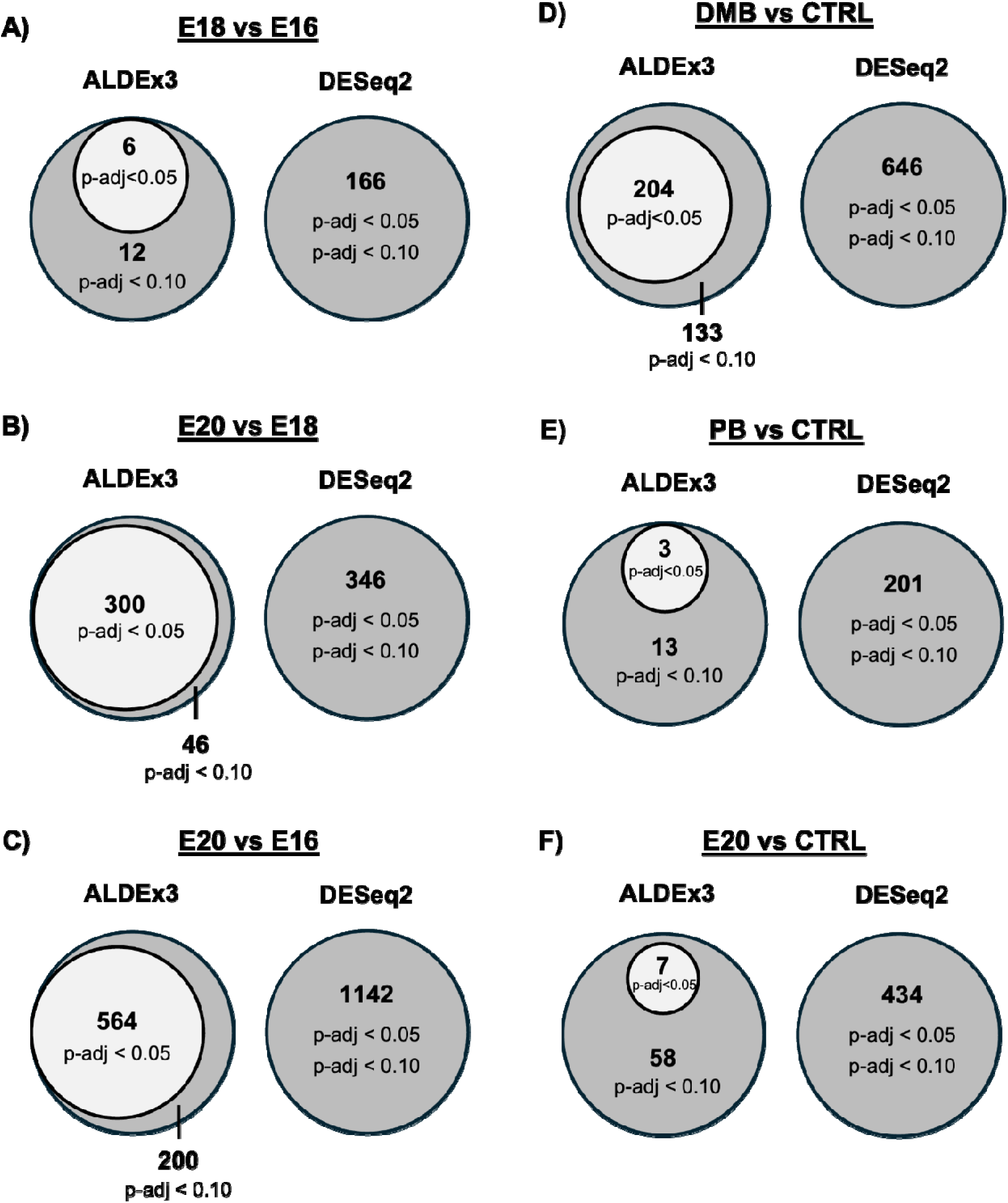
Significance threshold comparison between ALDEx3 and DESeq2. Venn diagrams comparing the number of DEGs identified at two significance thresholds (adj. p < 0.05, inner circle; adj. p < 0.10, outer circle) by ALDEx3 and DESeq2 for **A-C)** Developmental comparisons, **D-F)** Paralysis comparisons.

**Supplementary Figure 3:**
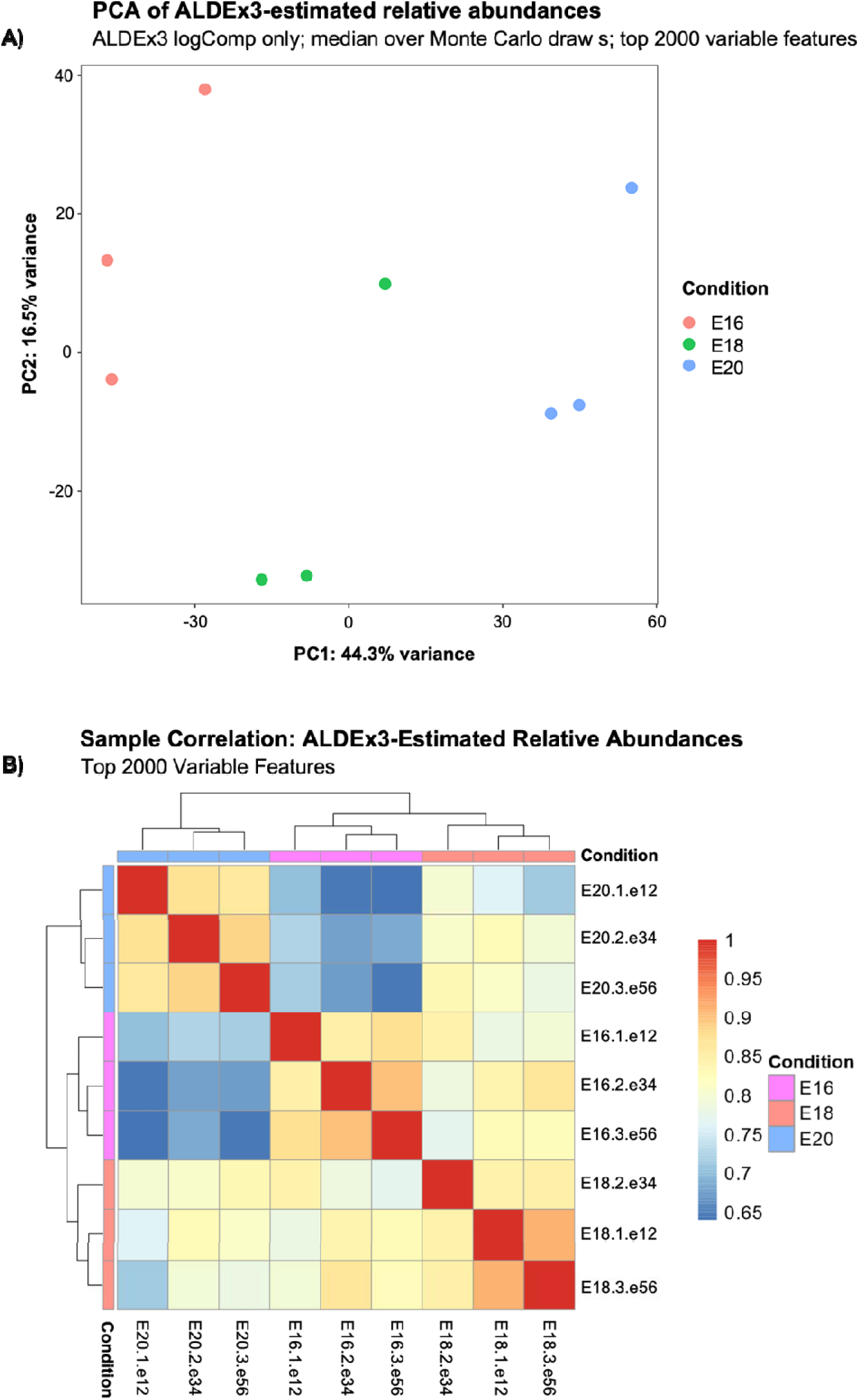
Sample clustering by ALDEx3-estimated relative abundances during development. **A)** PCA of ALDEx3 log-ratio-transformed relative abundances (median over Monte Carlo draws) for the top 2000 most variable features, colored by developmental timepoint. **B)** Hierarchically clustered heatmap of pairwise sample correlations based on the same top 2000 variable features. Color scale represents Pearson correlation coefficient.

**Supplementary Figure 4:**
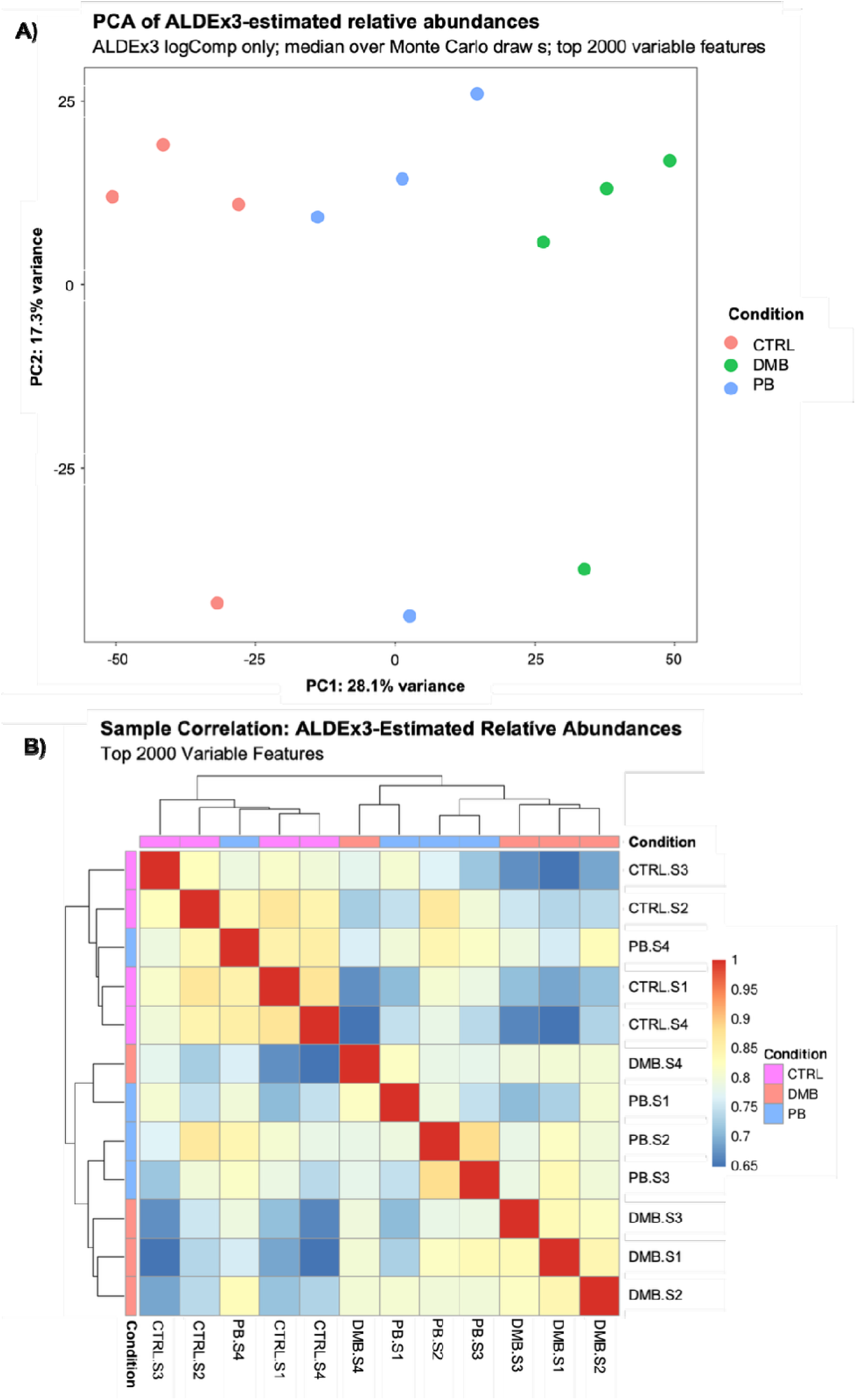
Sample clustering by ALDEx3-estimated relative abundances during paralysis. **A)** PCA of ALDEx3 log-ratio-transformed relative abundances (median over Monte Carlo draws) for the top 2000 most variable features, colored by paralysis treatment. **B)** Hierarchically clustered heatmap of pairwise sample correlations based on the same top 2000 variable features. Color scale represents Pearson correlation coefficient.

**Supplementary Figure 5:**
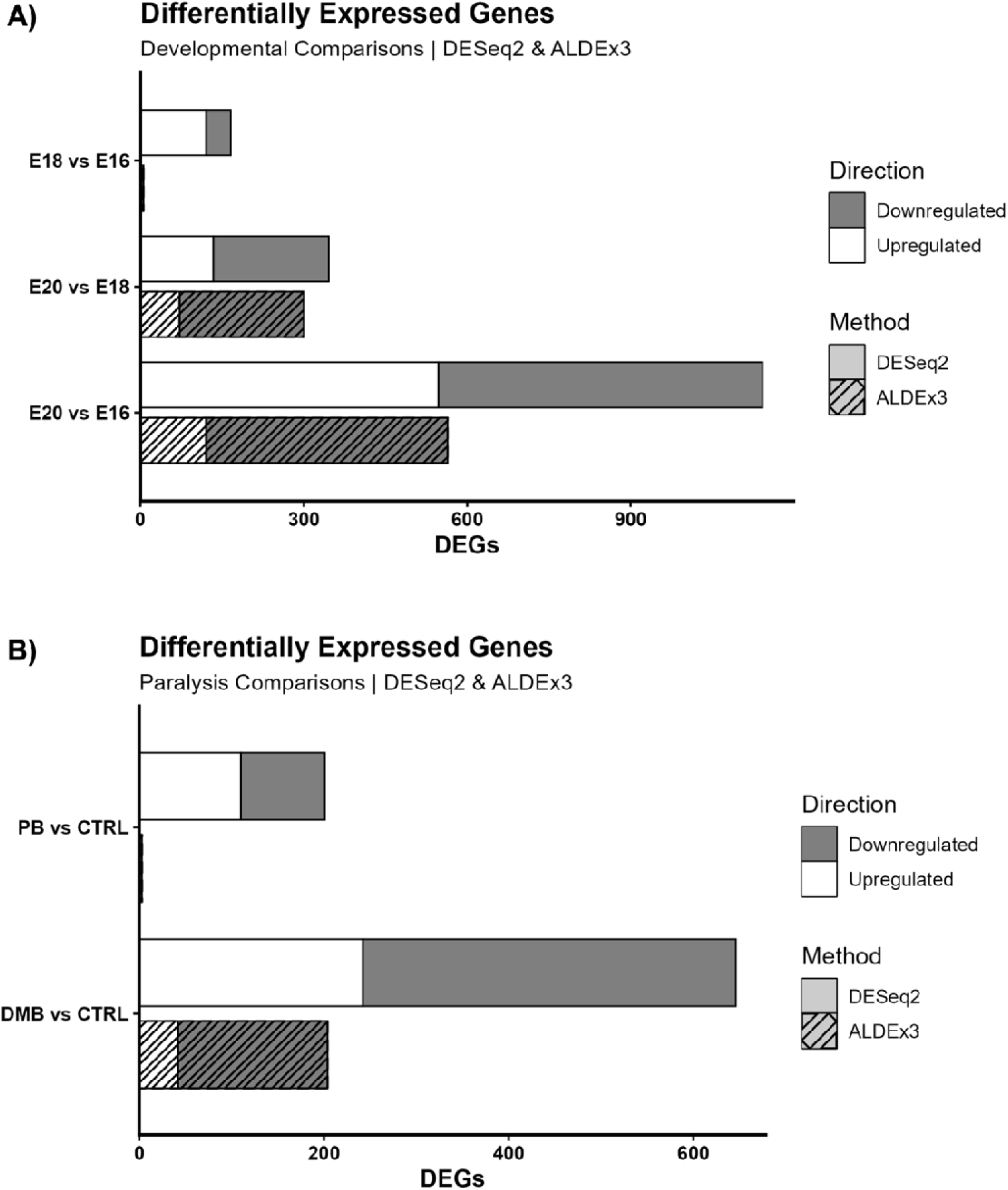
Directionality of differentially expressed genes. Bar plots showing the number of upregulated and downregulated DEGs identified by DESeq2 and ALDEx3 for **A)** developmental comparisons and **B)** paralysis comparisons. Bar shading indicates direction of expression change.

**Supplementary Figure 6:**
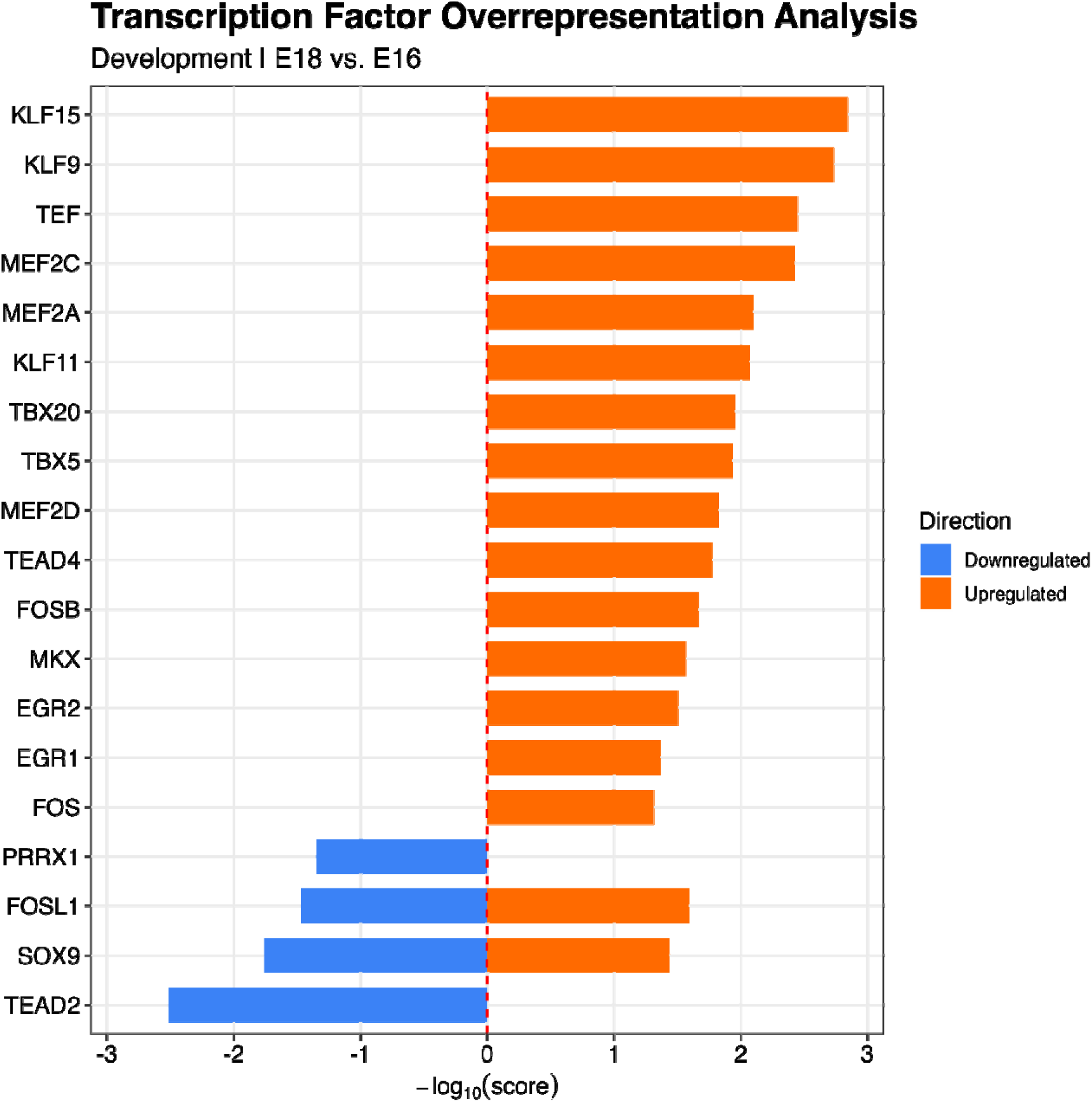
Transcription factor overrepresentation analysis, E18 vs. E16. Bar plot of significant transcription factors identified by ChEA3 (integrated score ≤ 0.05) among upregulated and downregulated DEGs for E18 vs. E16, ranked by −log10(score). Upregulated and downregulated gene sets were analyzed separately to account for directionality.

**Supplementary Figure 7:**
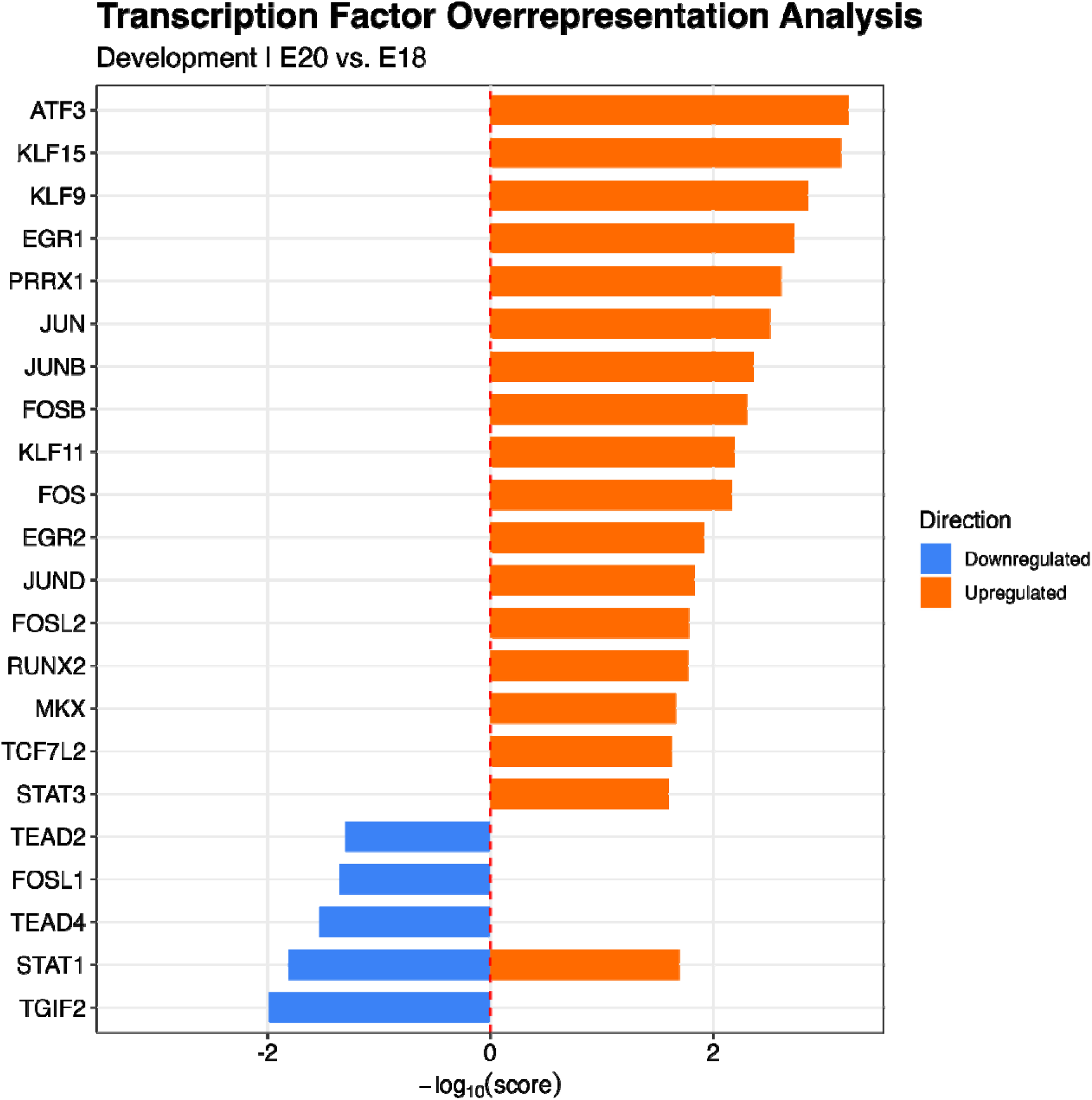
Transcription factor overrepresentation analysis, E20 vs. E18. Bar plot of significant transcription factors identified by ChEA3 (integrated score ≤ 0.05) among upregulated and downregulated DEGs for E20 vs. E18, ranked by −log10(score). Upregulated and downregulated gene sets were analyzed separately to account for directionality.

**Supplementary Figure 8:**
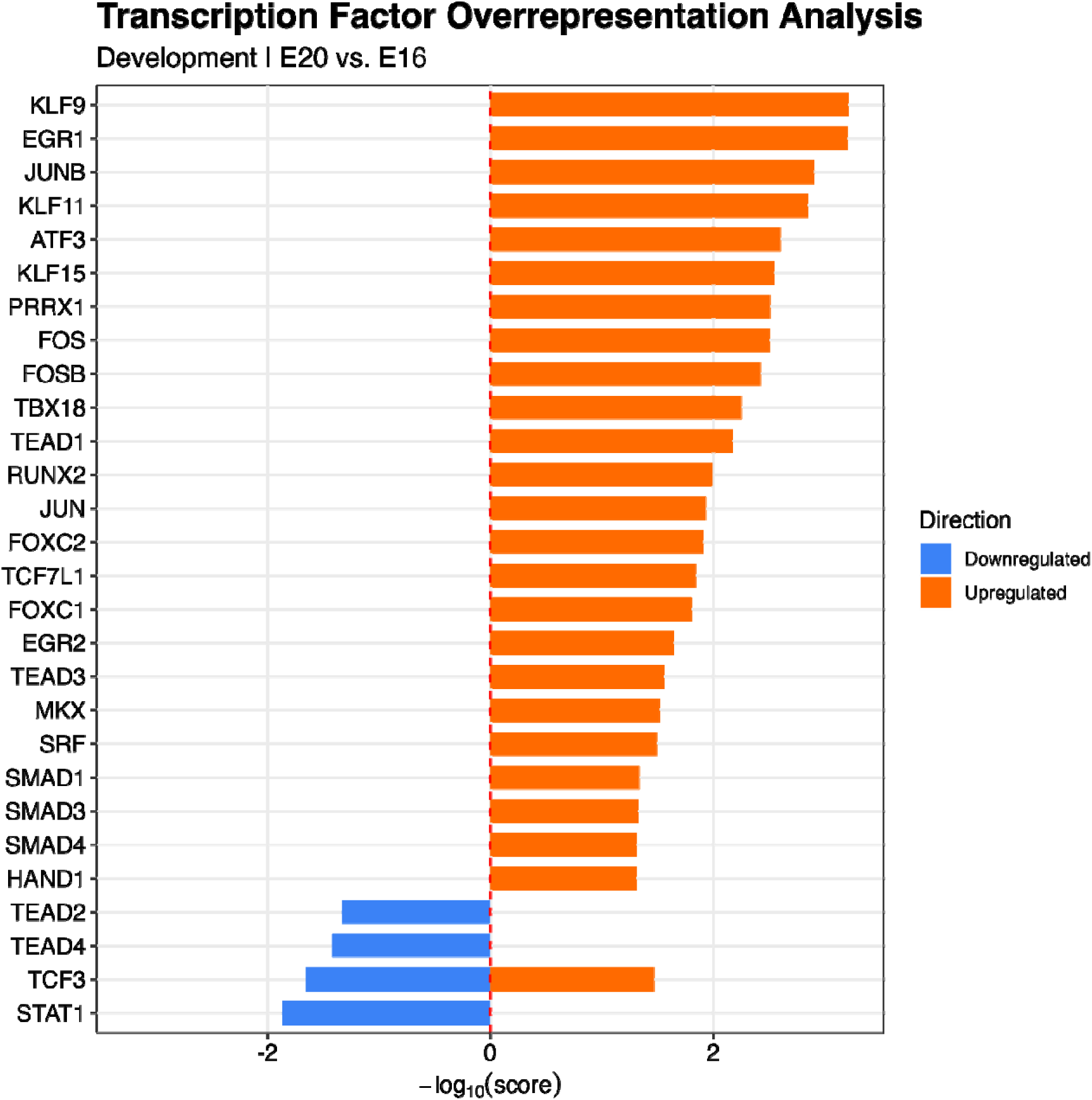
Transcription factor overrepresentation analysis, E20 vs. E16. Bar plot of significant transcription factors identified by ChEA3 (integrated score ≤ 0.05) among upregulated and downregulated DEGs for E20 vs. E16, ranked by −log10(score). Upregulated and downregulated gene sets were analyzed separately to account for directionality.

**Supplementary Figure 9:**
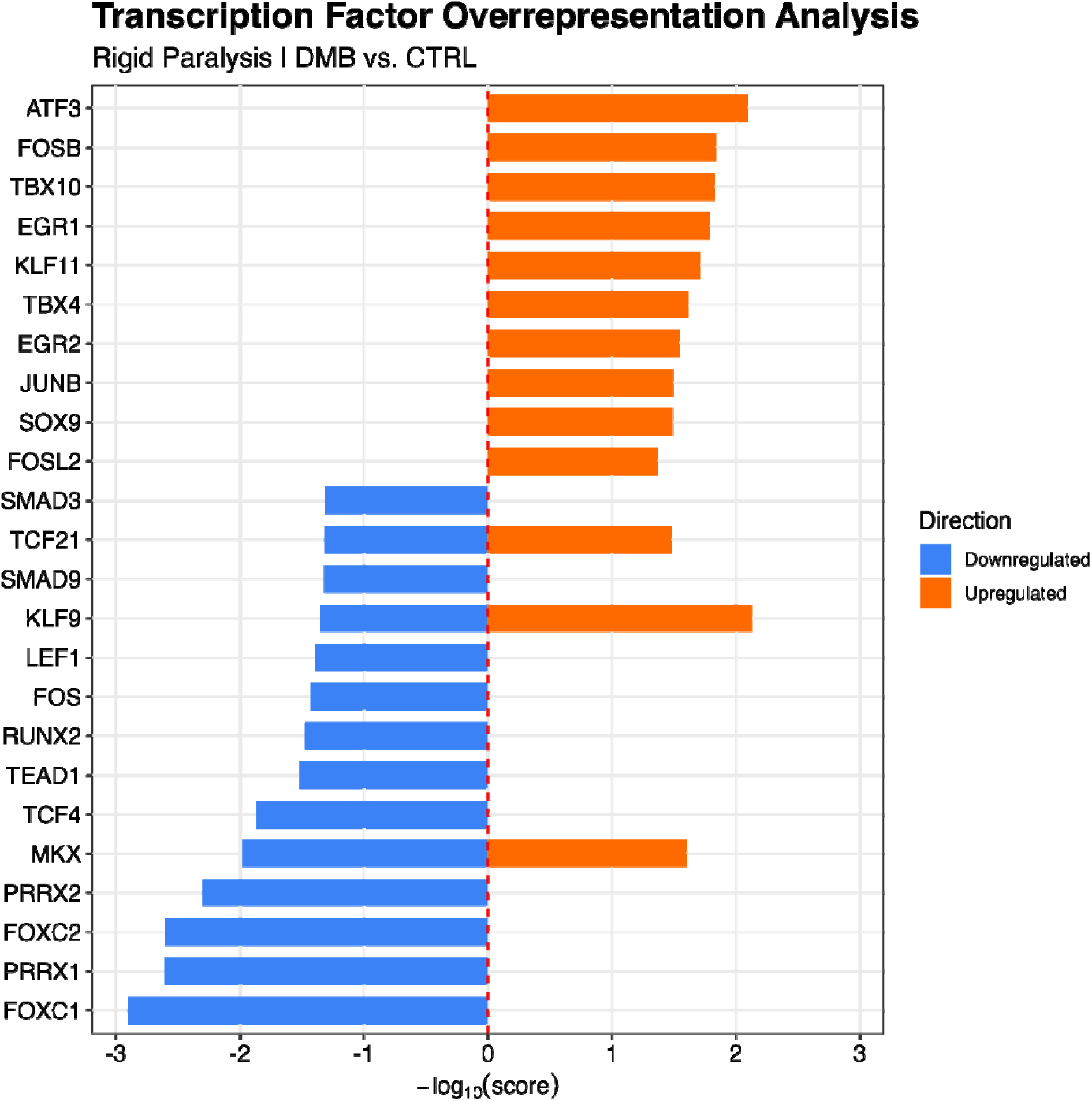
Transcription factor overrepresentation analysis, DMB vs. CTRL. Bar plot of significant transcription factors identified by ChEA3 (integrated score ≤ 0.05) among upregulated an downregulated DEGs for rigid paralysis ranked by −log10(score). Upregulated and downregulated gene sets were analyzed separately to account for directionality.

**Supplementary Figure 10:**
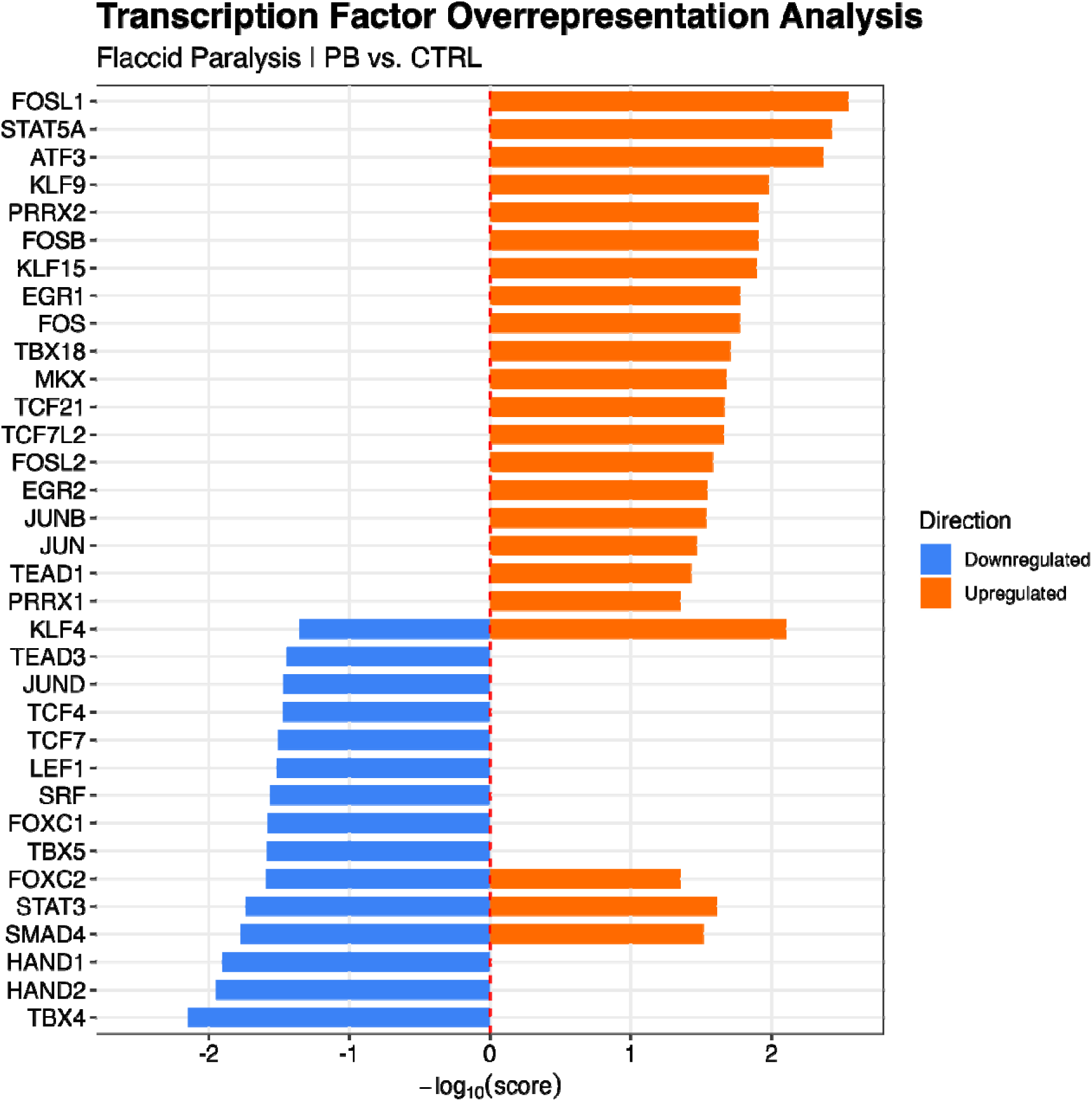
Transcription factor overrepresentation analysis, PB vs. CTRL. Bar plot of significant transcription factors identified by ChEA3 (integrated score ≤ 0.05) among upregulated an downregulated DEGs for flaccid paralysis ranked by −log10(score). Upregulated and downregulated gene sets were analyzed separately to account for directionality.

## Supplementary Tables

**Supplementary Table 1:** Sign-Flip analysis of differentially expressed genes across methods. Log2 fold-change sign flips (genes estimated to be upregulated by one method but downregulated by the other) are categorized by significance (adj. p-value < 0.05). “Significant in Both” represents sign-flips significant in both ALDEx3 and DESeq2 datasets. “Significant in DESeq2” represent sign-flips significant in the DESeq2 dataset. “All Genes” represents all sign-flips identified in our comparisons, regardless of statistical significance. Gene names are displayed in parentheses.

|  |  | Sign Flip |  |  |
|---|---|---|---|---|
| **Comparison** | **Figure** | **Significant in Both Methods** | **Significant in DESeq2** | **All DEGs** |
| E18 vs. E16 | 1D | 0 out of 4 | 1 out of 166 (THEM4) | 953 out of 11,741 |
| E20 vs. E18 | 1E | 1 out of 233 (IFNLR1) | 2 out of 356 (IFNLR1 and H2AC1) | 1,763 out of 11,741 |
| E20 vs. E16 | 1F | 0 out of 512 | 1 out of 1,142 (OLR1) | 1,744 out of 11,774 |
| DMB vs. CTRL | 2B | 0 out of 201 | 0 out of 646 | 964 out of 11,923 |
| PB vs. CTRL | 2D | 0 out of 3 | 0 out of 201 | 1,173 out of 11,923 |

